# Global genome analysis identifies glycolipids and lipoteichoic acid alanylation as contributors to Group B streptococcal diabetic wound infection

**DOI:** 10.1101/2025.05.02.651901

**Authors:** Luke R. Joyce, Madeline S. Akbari, Dustin T. Nguyen, Brady L. Spencer, Jeremy Rahkola, Kevin S. McIver, Alexander R. Horswill, Kelly S. Doran, Rebecca A. Keogh

## Abstract

Individuals with diabetes frequently develop chronic, hyper-inflammatory, non-healing wounds, which are the leading cause of all non-traumatic amputations. Group B *Streptococcus* (GBS) is a prominent bacterium isolated from diabetic wound infections and in a murine model of diabetic wound infection, GBS stimulates an influx of neutrophils into the wound. Utilizing a transposon mutant screen, we identified 291 bacterial genes required for fitness during diabetic wound infection including enzymes involved in glycolipid biosynthesis and lipoteichoic acid (LTA) alanylation. GBS mutants lacking either LTA alanylation (Δ*dltA*) or all glycolipids (Δ*iagB*) are attenuated in a murine diabetic wound infection. GBS induces primary and secondary degranulation in primary human neutrophils and the Δ*iagB* mutant is significantly more susceptible to neutrophil killing by cationic antimicrobial peptides and reactive oxygen species than Δ*dltA*. Finally, we found that depletion of neutrophils led to significantly greater bacterial recovery, highlighting the importance of neutrophil defense during diabetic wound infection.

## INTRODUCTION

Diabetes is a group of metabolic diseases, projected to affect over 700 million people by 2045 ^1^. A frequent complication of diabetes is the development of wounds on the foot and lower limbs ^2,3^. These wounds often become chronic, failing to heal within 4 to 6 weeks, due to an unresolved inflammatory phase ^4,5^. This inflammation, primarily driven by neutrophils and macrophages leads to the production of reactive oxygen species (ROS), cationic antimicrobial peptides (CAMPs), matrix metalloproteases, and pro-inflammatory chemokines and cytokines, which perpetuate inflammation and limit the reformation of tissue ^6,7^.

Diabetic wounds frequently become colonized by a variety of bacterial species which can result in chronic infection ^8–11^. While some infections are limited to the wound itself, if the infection is uncontrolled, bacteria can penetrate deeper tissue leading to bone and systemic infections, which drive the need for amputation ^3,12,13^. Some of the most prominent bacterial species identified in diabetic wound infection are *Staphylococcus aureus*, *Enterococcus faecalis*, Group B *Streptococcus* (GBS), *Pseudomonas aeruginosa* and many anaerobic species ^8,9,14,15^. These infections are often difficult to treat due to high levels of antibiotic resistance and tolerance amongst isolates ^8,16,17^. Interestingly, work by Sook et al. has demonstrated that *S. aureus* evolves antibiotic resistance in a diabetic host, and that hyperglycemia potentiates the expansion of resistant populations *in vivo* ^18^. Further complicating treatment of diabetic wounds is that bacteria in chronic wounds often live in biofilm communities ^19,20^. Biofilms, consisting of bacterial cells, DNA, and proteins often lead to a physical barrier which is difficult to treat by antibiotics or routine debridement of tissues ^19^. These biofilms can be mono or polymicrobial in nature, and work by Ch’ng et al. has found that *S. aureus* can assist in *E. faecalis* biofilm formation via heme cross-feeding ^21^.

GBS is a Gram-positive bacterium which colonizes approximately 25% of all individuals in the gastrointestinal and vaginal tract ^22–24^. In a healthy host, GBS typically colonizes without any adverse health outcomes; however, GBS can cause urinary tract infections ^25^ and during pregnancy, GBS can ascend vertically and cause intrauterine infections, leading to inflammation and adverse pregnancy outcomes ^26,27^. Further, gestational diabetes mellitus has been identified as a risk factor for increased GBS vaginal colonization ^28,29^. In immunocompromised individuals such as neonates, and adults with underlying conditions such as diabetes and cancer, GBS can cause diverse infections including pneumonia, meningitis, cardiovascular disease and sepsis ^30–34^. GBS is frequently isolated from diabetic wounds, despite it rarely colonizing the wounds of non-diabetic individuals ^11,14,15,35–40^. In addition, GBS alone is sufficient to cause infection and inflammation in the diabetic wound environment ^41^. Despite this, little is known about which bacterial factors aid in GBS wound establishment, immune cell evasion, and persistence in this niche.

To address this, we previously developed a murine model of GBS diabetic wound infection and found that GBS infection is exacerbated in a diabetic host when compared to a non-diabetic ^41^. RNA-seq on wound tissues revealed that diabetic mice upregulate numerous genes encoding neutrophil effectors such as MPO, elastase, the subunits of calprotectin, as well as pro-inflammatory cytokines and chemokines in response to GBS infection when compared to uninfected controls. We further found that the abundance of these neutrophil effectors was significantly higher in diabetic mice in comparison to non-diabetic upon GBS infection. These data support that GBS encounters high levels of inflammation in the diabetic wound microenvironment, but currently, little is known regarding the bacterial factors that promote survival in this niche. To date only the hemolysin/cytolysin (β-H/C, hemolytic pigment) and the plasminogen binding protein, PbsP, have been shown to contribute to GBS diabetic wound infection ^41^.

Here, we observe that GBS infection drives an influx of neutrophils into the diabetic wound microenvironment, and that GBS and neutrophils co-localize spatially in diabetic wound tissues. We next utilize a global unbiased approach of transposon sequencing (Tn-seq) and identify GBS genes that are required for fitness in the diabetic wound environment. We find 291 genes that are significantly underrepresented in this niche, including genes involved in metabolism, capsule biosynthesis, the production of membrane glycolipids and lipoteichoic acid (LTA) alanylation. Membrane lipids are a vital part of host-pathogen interactions ^42^ and we have recently characterized the role of GBS glycolipids during systemic infection and meningitis disease progression ^43–45^. Utilizing GBS glycolipid mutants and a strain deficient in LTA alanylation, we confirm our Tn-seq results by finding that all four mutants are attenuated in diabetic wound infection. We further show the importance of GBS glycolipids and LTA alanylation to keratinocyte interactions and survival against neutrophil killing by cationic antimicrobial peptides (CAMPs) and reactive oxygen species (ROS). Together, these data demonstrate the global gene requirement for GBS fitness in the diabetic wound and further identify GBS glycolipids and LTA alanylation as contributors to neutrophil survival in this niche.

## RESULTS

### Neutrophils are the most abundant cell type GBS interacts with during diabetic wound infection

The diabetic wound is a hyper inflammatory niche with an abundance of immune cells such as phagocytes present ^6,46^, however, it is unknown which immune cell populations are the most abundant in response to GBS infection. To address this, we wounded diabetic mice and inoculated the wounds with either GBS WT or PBS controls (*n* = 7 per group) and at 48 hpi, euthanized and perfused mice before harvesting wound tissues. We used flow cytometry to quantify different populations of immune cells ^47^ (Figure S1A). The primary populations of immune cells present in both the uninfected (mock) and GBS infected wounds were neutrophils, followed by macrophages (Figure 1A). Upon infection with GBS, we observed a significant decrease in the numbers of live cells and live CD45^+^ cells as a percent of collected singlets (Figure S1B,C), suggesting greater cell death, inclusive of immune cells, in the infected wounds compared to uninfected controls. Live neutrophils as a percent of collected singlets did not change between uninfected and infected wounds (Figure S1D); however, neutrophils did represent a greater proportion of immune cells as the percent of neutrophils of live CD45^+^ cells compared to the mock controls (Figure 1B). Additionally, a significant decrease in the percent of macrophages of live CD45^+^ cells compared to mock controls was observed (Figure 1B). This suggests that despite increased immune cell death, the remaining live immune cell population is skewed further toward neutrophils in infected wounds. This is consistent with our previous RNA sequencing results that demonstrated a greater abundance of neutrophil effectors such as MPO and elastase, calprotectin, and CXCL1 in GBS infected diabetic wound tissues when compared to uninfected mice ^41^. No differences in the CD11b^-^CD11c^-^ (including lymphocytes) and the MHCII^-^SSC^lo^ (including monocytes and NK cells) populations as a percent of CD45^+^ cells were observed (Figure S1E,F). A significant decrease in the percent of CD24^+^ cells and dendritic cells was observed (Figure S1G,H), possibly due to the proportional increase of neutrophils in the wound. Additionally, we did not observe major alterations in percentages of immune cell populations in the blood of GBS infected and uninfected controls (Figure S1I). These data indicate that neutrophils are the most abundant immune cell type GBS encounters in the wound microenvironment at this time point of infection.

**Figure 1.**
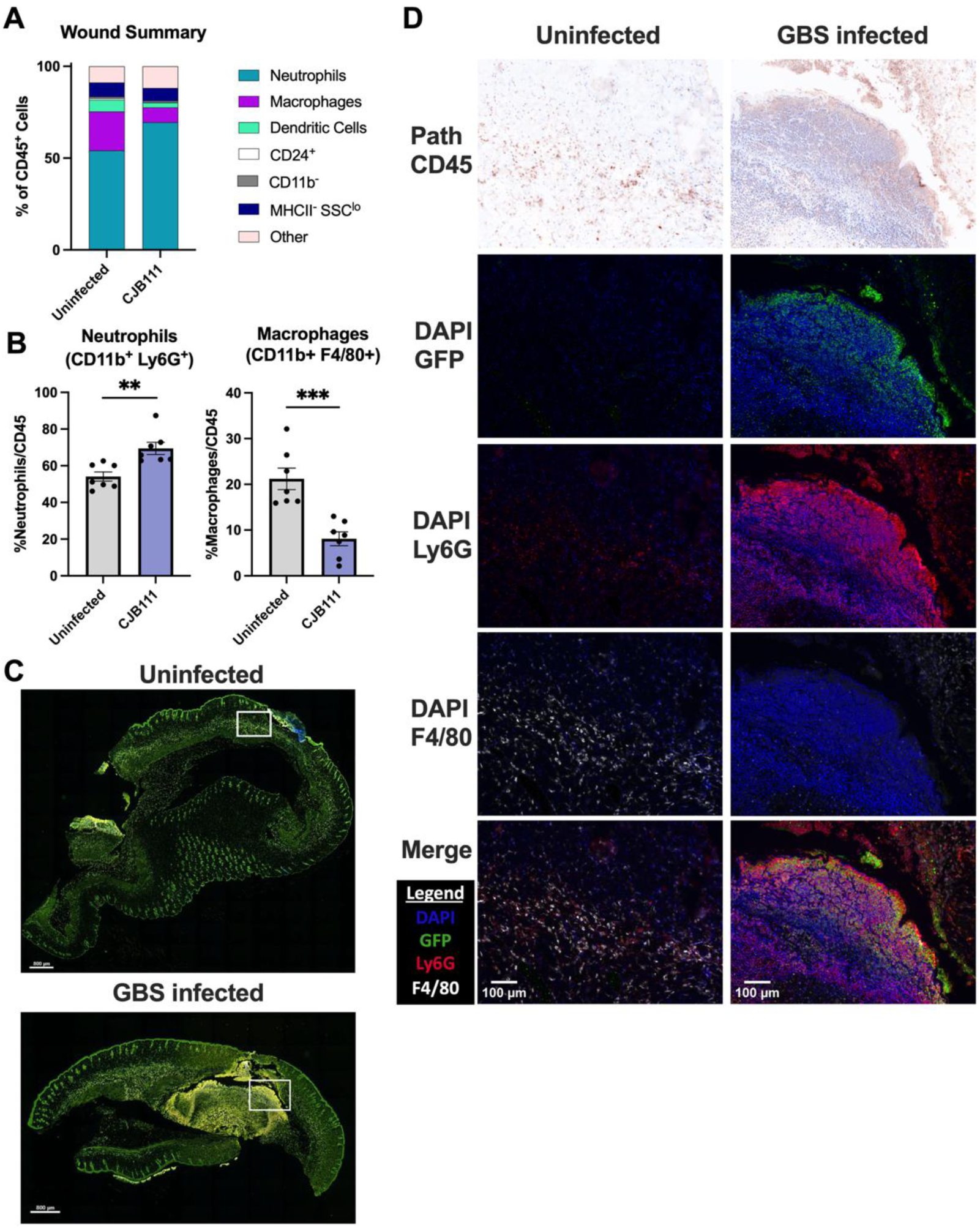
Neutrophils are abundant in the diabetic wound during infection. A) Wound immune cell populations measured by flow cytometry. B) A significant increase of neutrophils (left) and significant decrease of macrophages (right) as a percent of CD45^+^ immune cells is observed in GBS infected diabetic wounds at 48 h compared to uninfected diabetic wounds. C) Multispectral immunohistochemistry microscopy images of diabetic wound tissues mock or GBS infected. Scale bar 800 µm. D) White boxed region of interested showing CD45 pathology, GFP expressing GBS in the wound, Ly6G^+^ neutrophil marker, F4/80^+^ macrophage marker, and merged image. Scale bar 100 µm. Significance determined by B) two-tailed T-Test; **p<.01; ***p<.001.

Next, we utilized the same innate immune flow panel to determine which immune cells were present in a non-diabetic vs. a diabetic host during GBS infection (Figure S2). Interestingly, we observed a significant decrease in live cells as percentage of singlets as well as a significant decrease in CD45^+^ cells as a proportion of live cells in a diabetic host when compared to non-diabetic. These data suggest that the non-diabetic host has more live immune cells in response to GBS infection in the wound (Figure S2A,B). Neutrophils were the most abundant population amongst live cells in both a non-diabetic and diabetic host, highlighting their importance in combatting GBS during wound infection regardless of diabetic status. However, there were significantly more neutrophils in a non-diabetic host than diabetic (Figure S2C,D), and no differences between the other immune populations (Figure S2E-J).

To further confirm the immune profile of diabetic wounds, we utilized multispectral quantitative imaging of wound tissues excised from diabetic mice infected with WT GBS fluorescently tagged with GFP or the mock PBS control. Infected mice developed a robust wound under the epidermal layer (Figure 1C) when compared to the uninfected control. We then chose a region of interest spanning the wound and epidermal layer (boxed in white) for further analysis of immune populations. Pathology of the GBS infected region of interest confirmed a greater proportion of immune cells when compared to the PBS control. GBS co-localized with Ly6G positive cells, confirming the influx of neutrophils to the site of infection (Figure 1D). Conversely, higher levels of F4/80 staining were seen in the PBS control group. These data are the first to demonstrate the spatial dynamics of GBS and immune cells during diabetic wound infection and highlight that GBS and neutrophils interact within this microenvironment.

### Genome-wide analysis of GBS factors required for survival in the diabetic wound

We next sought to identify which genetic factors GBS requires to colonize and survive in the diabetic wound. We utilized a GBS Tn mutant library ^48–50^ in the CJB111 (Serotype V) background ^51,52^ and our previously described diabetic wound infection model as described in materials and methods ^41^. Groups of mice (4 per group) were infected with ∼1.5 x 10^7^ CFU of the GBS Tn library in biological triplicate. Mice were monitored over the course of infection, and at 48 hours post infection (hpi) mice were euthanized and wound tissue collected (Figure 2A). The input and wound recovered libraries were processed as described in the materials and methods.

**Figure 2.**
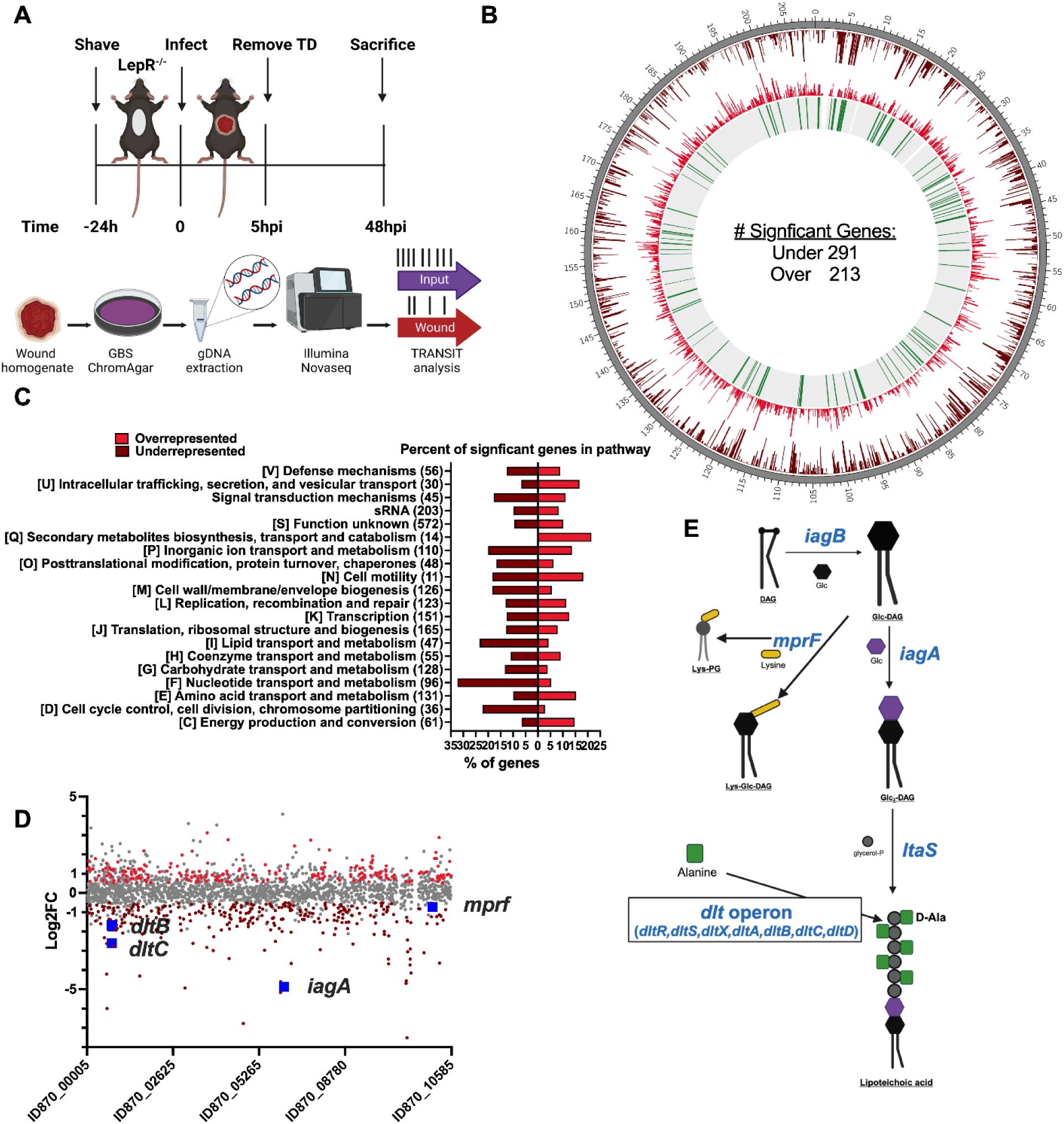
Essential GBS Genes During Diabetic Wound Infection. A) Experimental set-up for Tn-seq. B) CIRCOS plot showing from outer ringer to inner ring; dark grey, genome length in Kb; maroon, negative Log2Fold Change (L2FC); red, positive L2FC; statistically significant genes (adj. p-value ≤ 0.05) shown in green and non-significant genes in light grey. C) COG pathway analysis of percent of statistically significant genes in each COG for both over- (red) and under- (maroon) represented genes. (Numbers) indicate the total number of genes in each pathway. D) Linear plot of significantly over (red) and underrepresented (maroon) genes ordered by genomic position. E) Glycolipid biosynthesis and LTA decoration pathway in GBS.

To identify transposon insertion sites, sequenced reads were mapped to the GBS CJB111 genome (CP063198) (Figure S3), which identified 291 genes as significantly underrepresented (adj. p-value < 0.05, log2FC < -0.5) and 213 genes as significantly overrepresented (adj. p-value < 0.05, log2FC > 0.5) in the wound compared to the input library (Figure 2B) (Table S1). The significant gene hits are distributed across the genome and were then assigned clusters of orthologous groups of proteins (COGs). The number of significant gene hits in each COG were normalized to the total number of genes in each COG, to determine the percentage of genes in each COG that were under- or overrepresented *in vivo*. COGs including nucleotide transport and metabolism, carbohydrate transport and metabolism, inorganic ion transport and metabolism, lipid transport and metabolism, and cell wall/membrane/envelope biogenesis had the greatest proportion of underrepresented genes in the diabetic wound (Figure 2C). We identified multiple genes known to contribute to GBS infection as significantly underrepresented such as genes that are involved in β-H/C biosynthesis, capsule biosynthesis, two-component regulatory systems, glutamine transport, and purine metabolism (Table 1 and Table S1).

**Table 1.**
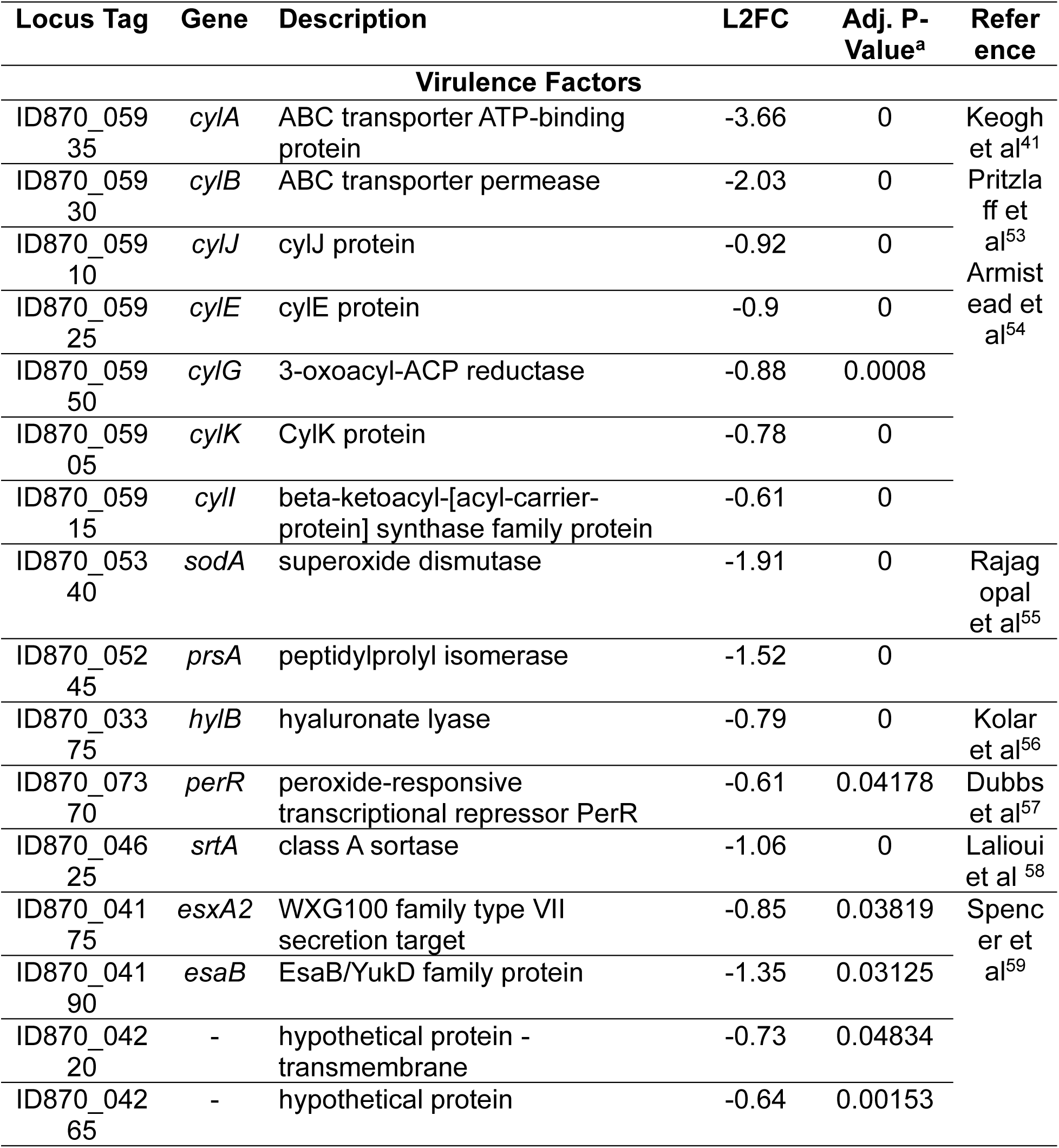
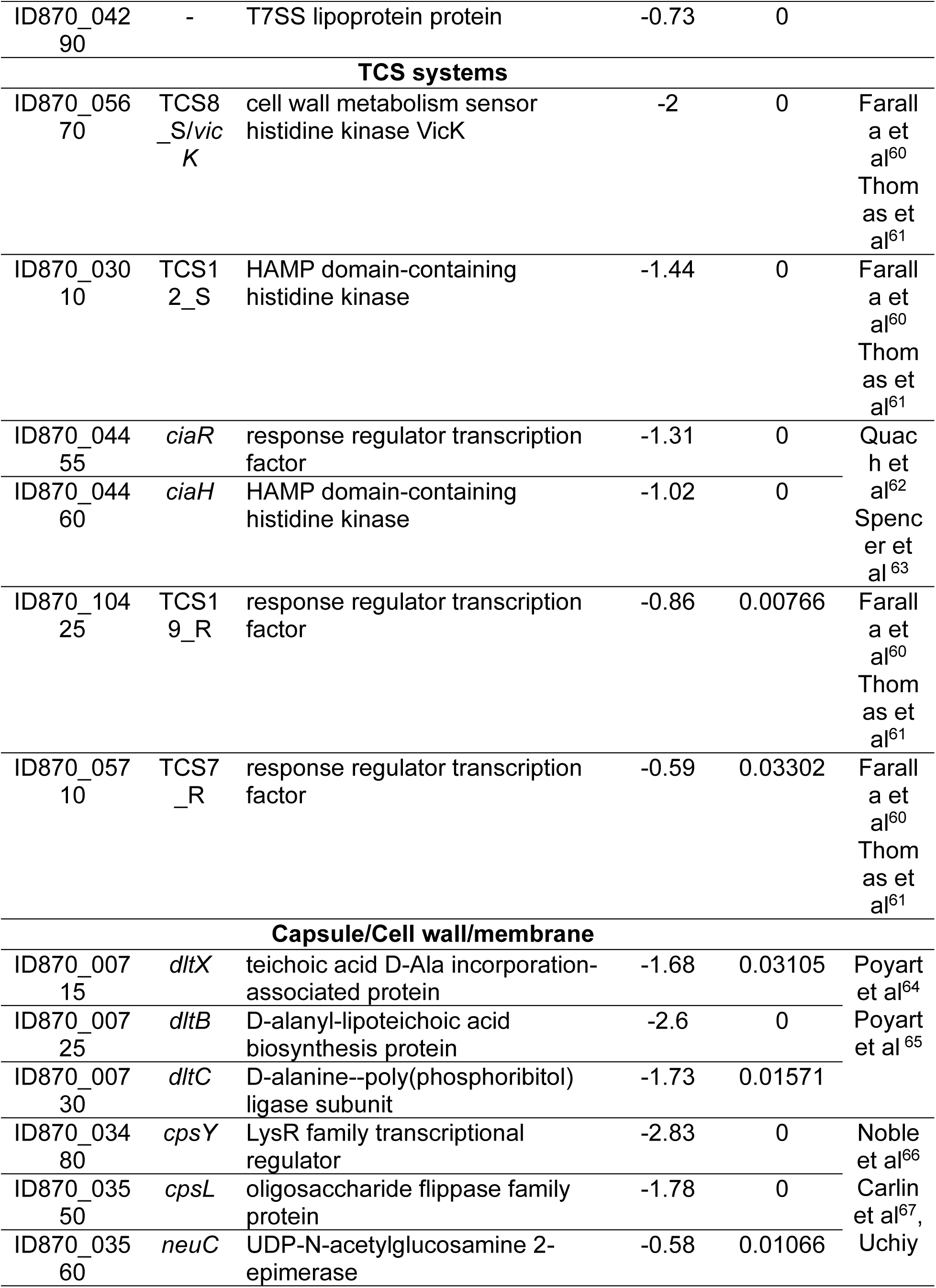
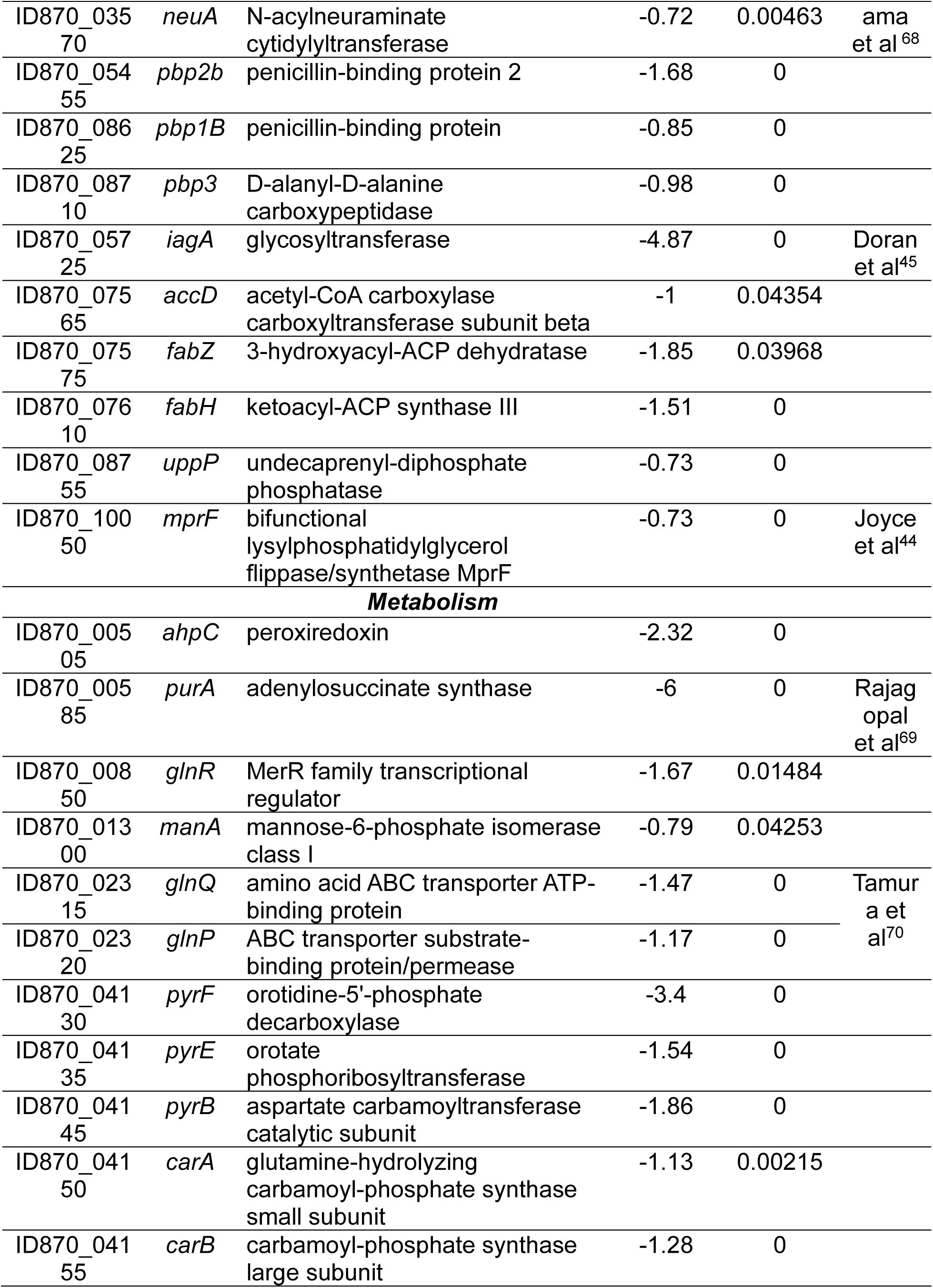
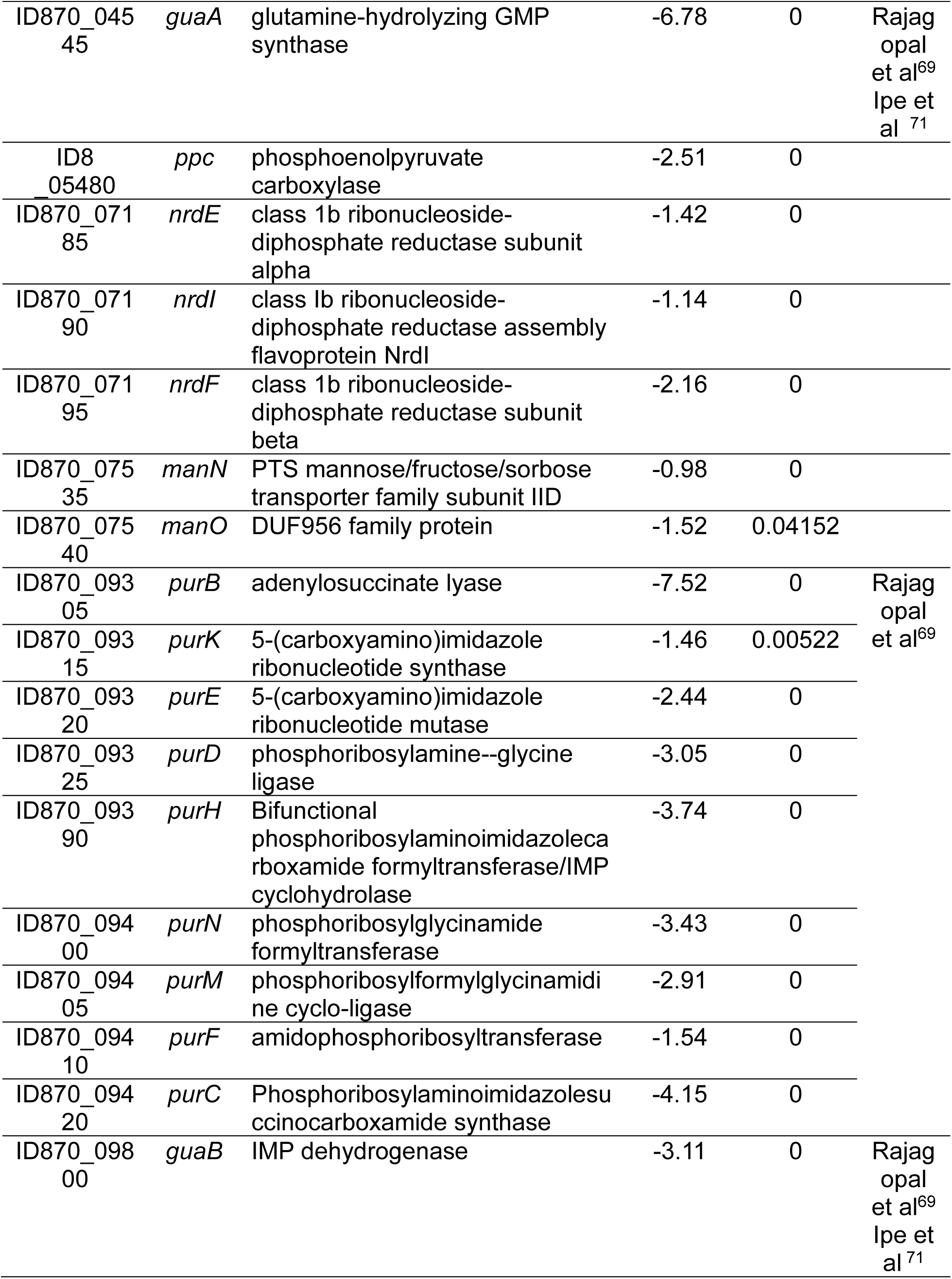
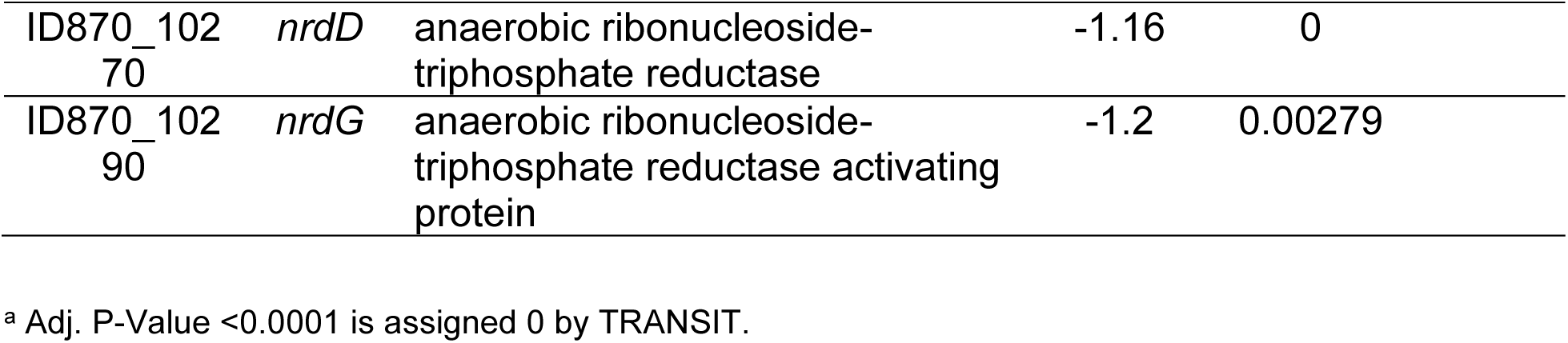
Significantly underrepresented genes in the diabetic wound.

Interestingly, among the most significantly underrepresented genes was *iagA* (Figure 2D, Table 1) which is a glycosyltransferase involved in the biosynthesis of the LTA anchor, diglucosyl-diacylglycerol (Glc_2_-DAG) in GBS ^43,45^ (Figure 2E), and is in an operon with the first glycolipid biosynthesis gene, *iagB*, that is responsible for glucosyl-DAG (Glc-DAG) biosynthesis ^43^. An additional gene in glycolipid biosynthesis pathway, *mprF* responsible for synthesizing the amino acylated glycolipid lysyl-glucosyl-DAG (Lys-Glc-DAG) ^44,72^ and multiple genes involved in the alanylation of LTA (*dlt* operon) ^65^ were also significantly underrepresented in the diabetic wound compared to input (Figure 2D,E). The two remaining genes in the biosynthetic pathway, namely *iagB* and *ltaS* genes, were not identified in the Tn-seq dataset. However, upon further investigation, we observed that there were very few to no transposon insertions present in these genes in the input libraries, thus making it hard to identify their importance with this approach. GBS glycolipids have recently been shown to impact GBS bloodstream survival ^43^ and disruption and inflammation of the blood-brain barrier (BBB) ^44,45^, while alanylation of LTA has been shown to be important for bloodstream survival ^64^; yet the contribution of these pathways to GBS survival in the diabetic wound have yet to be elucidated.

### GBS glycolipids and LTA alanylation are required for epithelial cell invasion and survival in the diabetic wound

To colonize the diabetic wound, GBS must interact with epidermal keratinocyte cells, a highly specialized epithelial cell ^73^. To investigate the direct interaction between GBS WT CJB111 as well as isogenic glycolipid mutants in *iagB*, ^43^, *iagA* ^43^, *mprF* ^44^ and *dltA* and human keratinocytes, we used the immortalized human keratinocyte cell line, N/TERT-2G ^74^. GBSΔ*iagB* is devoid of all glycolipids, GBSΔ*iagA* lacks Glc_2_-DAG the LTA anchor, GBSΔ*mprF* lacks the lysine decorated lipids Lys-Glc-DAG and Lys-PG, and the GBSΔ*dltA* mutant is unable to add alanine modifications to its LTA polymer ^65^ (Figure 2E). *In vitro* assays for adhesion and invasion were performed as described previously for other cell lines ^63,75^. There was no significant difference in the ability of the mutant strains to adhere to N/TERT cells (Figure 3A); however, there were significantly fewer mutant bacteria recovered from the intracellular compartment compared to GBS WT (Figure 3B) indicating GBS glycolipids and LTA alanylation may play roles in invasion of human keratinocytes. Furthermore, it was observed that GBSΔ*iagB* has significantly reduced intracellular survival over time compared to GBS WT, whereas all other mutant strains survived like GBS WT (Figure 3C). These observations indicate GBS glycolipids and LTA alanylation may contribute to GBS persistence in the wound.

**Figure 3.**
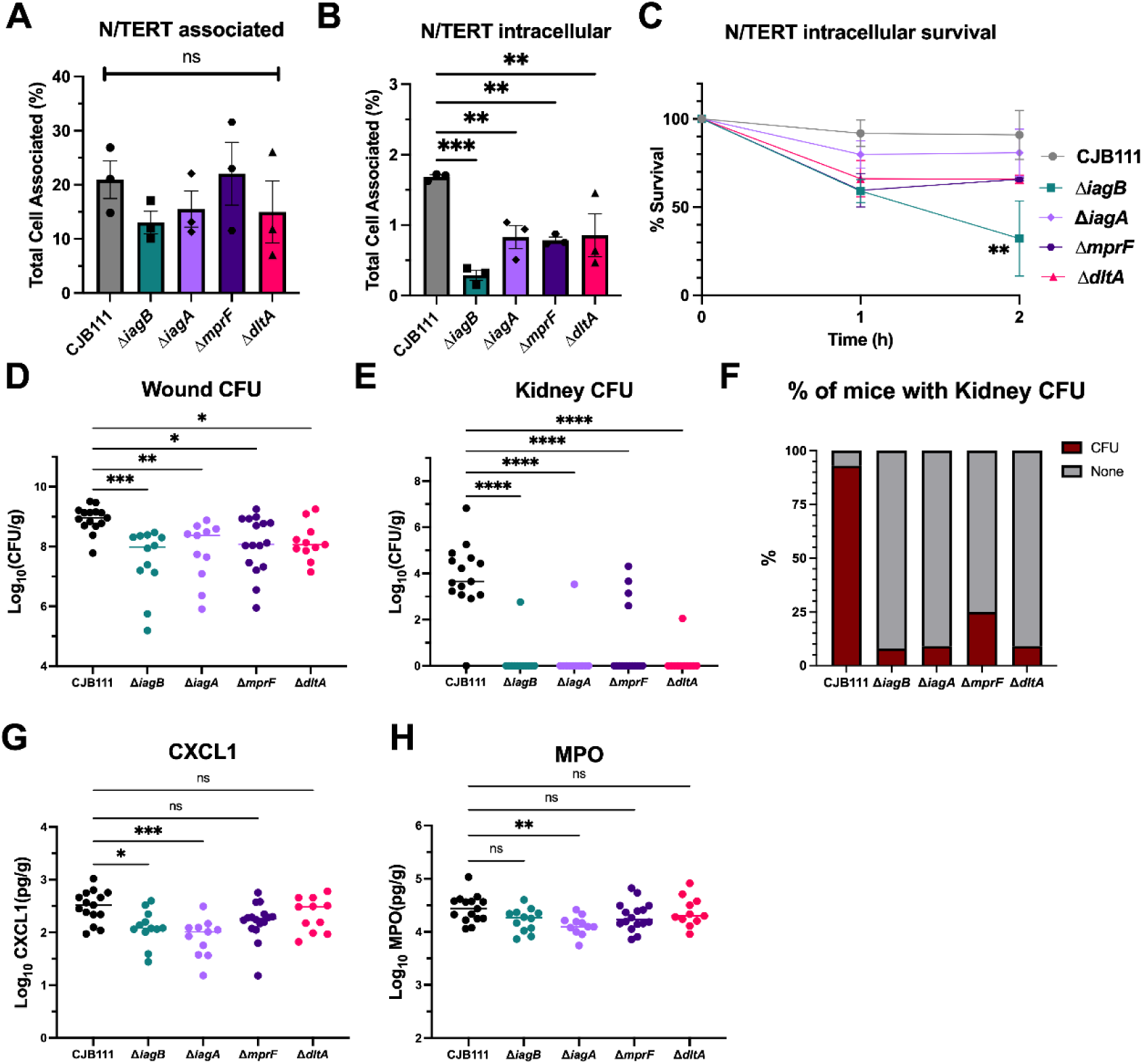
Glycolipid biosynthesis enzymes and LTA alanylation are necessary for epithelial cell invasion and survival in diabetic wound infection. *In vitro* assays for A) total cell associated (adherence) and B) intracellular GBS after 2 h infection, of GBS mutants N/TERTs indicate GBS glycolipids and LTA alanylation are dispensable for adhesion but are important for invasion of skin epithelial cells. C) Intracellular survival of GBS mutants over 2 h indicates GBS glycolipids are important for intracellular survival. Each assay was performed in biological triplicate, with three technical replicates per assay. Mean and SEM indicated. Diabetic mice were inoculated with 1 x 10^7^ CFU of GBS WT, Δ*iagB*, Δ*iagA*, Δ*mprF*, and Δ*dltA* isogeneic mutants and bacterial burdens were assessed after 48 h in the D) wound and E) kidneys. Significantly less CFU of all four mutants are recovered from the wound and kidneys of infected mice. F) % of mice in each infection group with detectable CFU in kidneys. G) Chemokine CXCL-1 and H) myeloperoxidase protein production measured by ELISA from wound tissues. Median indicated. GBS WT; *n =* 15, Δ*iagB; n* = 12, Δ*iagA*; *n* = 11, Δ*mprF*; *n* = 16, and Δ*dltA;* n *=* 11. Significance determined by A-C) One-way ANOVA with Fisher LSD and D,E,G,H) Kruskal-Wallis with Dunn’s multiple comparisons test; *p<.05, **p<.01, ***p<.001, ****p<.0001. ns; not significant.

To assess if there were alterations to the cell envelope of the mutants that may impact their interactions with keratinocytes, we performed transmission electron microscopy (TEM) (Figure S4A). We observed that GBSΔ*iagB* appears to have a general thickening of the cell envelope, along with GBSΔ*mprF* appearing to have minor thickening as well. GBSΔ*iagA* has no difference in appearance compared to GBS WT. Interestingly, GBSΔ*dltA* has a significantly altered inner electron dense ring compared to GBS WT and the other mutants, confirming prior work by Saar-Dover *et al*.^76^ In general all strains retained the polysaccharide capsule as assessed by flow cytometry using anti-sera targeting the type V capsule (Figure S4B). Interestingly, both GBSΔ*iagB* and GBSΔ*iagA* exhibited a slight, but significant, increase in capsule abundance compared to WT, which was restored to WT levels in the complemented strains. Additionally, no difference in membrane fluidity was observed between all strains as measured by anisotropy (Figure S4C).

To confirm our Tn-seq results and to elucidate the role of glycolipids and LTA alanylation during diabetic wound infection, we utilized our established streptozotocin induced diabetic wound model of infection ^77^ with GBS WT CJB111, Δ*iagB*, Δ*iagA*, Δ*mprF* and *dltA* mutant strains. After 48 hpi, all mutants were significantly attenuated in comparison to WT GBS (Figure 3D). Notably, in addition to having reduced bacterial burden in the wound, mutant strains were not able to spread systemically compared to WT GBS as evidenced by significantly reduced recovery of mutant bacteria from the kidneys (Figure 3E). Additionally, far fewer mice had detectable CFU in the kidneys compared to WT infected mice (Figure 3F). These data indicate that GBS glycolipids and LTA alanylation play an important role in GBS wound survival and systemic spread during diabetic infection.

Next, we assessed how GBS glycolipids and LTA modification would impact the abundance of neutrophil chemoattractant, CXCL1, and the leukocyte-derived enzyme myeloperoxidase (MPO) in the diabetic wound at the endpoint of infection. Wounds from mice infected with GBSΔ*iagB* and GBSΔ*iagA* had a significant decrease in both CXCL1 compared to GBS WT whereas wounds infected with GBSΔ*mprF* and GBSΔ*dltA* had no significant difference in CXCL1 abundance (Figure 3F). Interestingly, only GBSΔ*iagA* infected wounds had a significant decrease in the abundance of MPO compared to GBS WT infected wounds, suggesting high levels of neutrophils were still present in the GBS*ΔiagB,* GBS***Δ****mprF,* and GBS*ΔdltA* wounds to combat the infection (Figure 3G). We next utilized an early time point (5 hpi), to determine whether the loss of glycolipids impacted the influx of neutrophils to the site of infection. Mice infected with either WT, GBSΔ*iagB* (to determine impact of all glycolipids) and GBSΔ*dltA,* showed similar CFU recovery and abundance of CXCL1 and MPO (Figure S5). These data suggest that WT and mutant strains encounter similar levels of neutrophils early in infection and that the enhanced susceptibility of the GBSΔ*iagB* and GBSΔ*dltA* mutants to neutrophil killing led to enhanced clearance by 48 h.

### GBS glycolipids and LTA alanylation are vital for surviving neutrophil killing

To further understand the interaction between GBS and neutrophils, we isolated primary human neutrophils from four independent donors (two female and two male) to assess primary human neutrophil killing of GBS in diabetic glucose levels. Primary human neutrophils were conditioned in media containing 22 mM glucose to mimic hyperglycemia ^46^ before being exposed to opsonized bacteria. To assess the role of all GBS glycolipids, including the LTA anchor, we used a GBSΔ*iagB* mutant and to assess impact of LTA alanylation we used the GBS*ΔdltA* mutant. A significant decrease in survival was observed for GBSΔ*iagB* compared to GBS WT, which is consistent to our previous study ^43^ of GBS*ΔiagB* susceptibility to neutrophil killing under non-diabetic conditions; however, no significant difference was observed between GBSΔ*dltA* and GBS WT (Figure 4A). The phenotype of the GBS*ΔiagB* mutant was rescued when the strain was complemented with a plasmid containing the *iagB* gene *(+*pJIagB). GBS*ΔdltA* is known to be more susceptible to neutrophil killing and bloodstream clearance ^64^ so we assessed opsonophagocytic neutrophil mediated killing over time using the human HL60-neutrophil like cell line that is routinely maintained and differentiated in high glucose media. Interestingly, we observed that GBS*ΔiagB* is significantly more susceptible to neutrophil killing at 1, 2 and 3 h compared to GBS WT, whereas only after 2 and 3 h of infection was there a significant difference between GBSΔ*dltA* and GBS WT (Figure 4B). Additionally, after 3 h of exposure there was less than 1% survival for both mutants whereas WT maintains approximately 15% survival. These data indicate that GBS glycolipids and LTA alanylation both contribute to thwarting neutrophil clearance.

**Figure 4.**
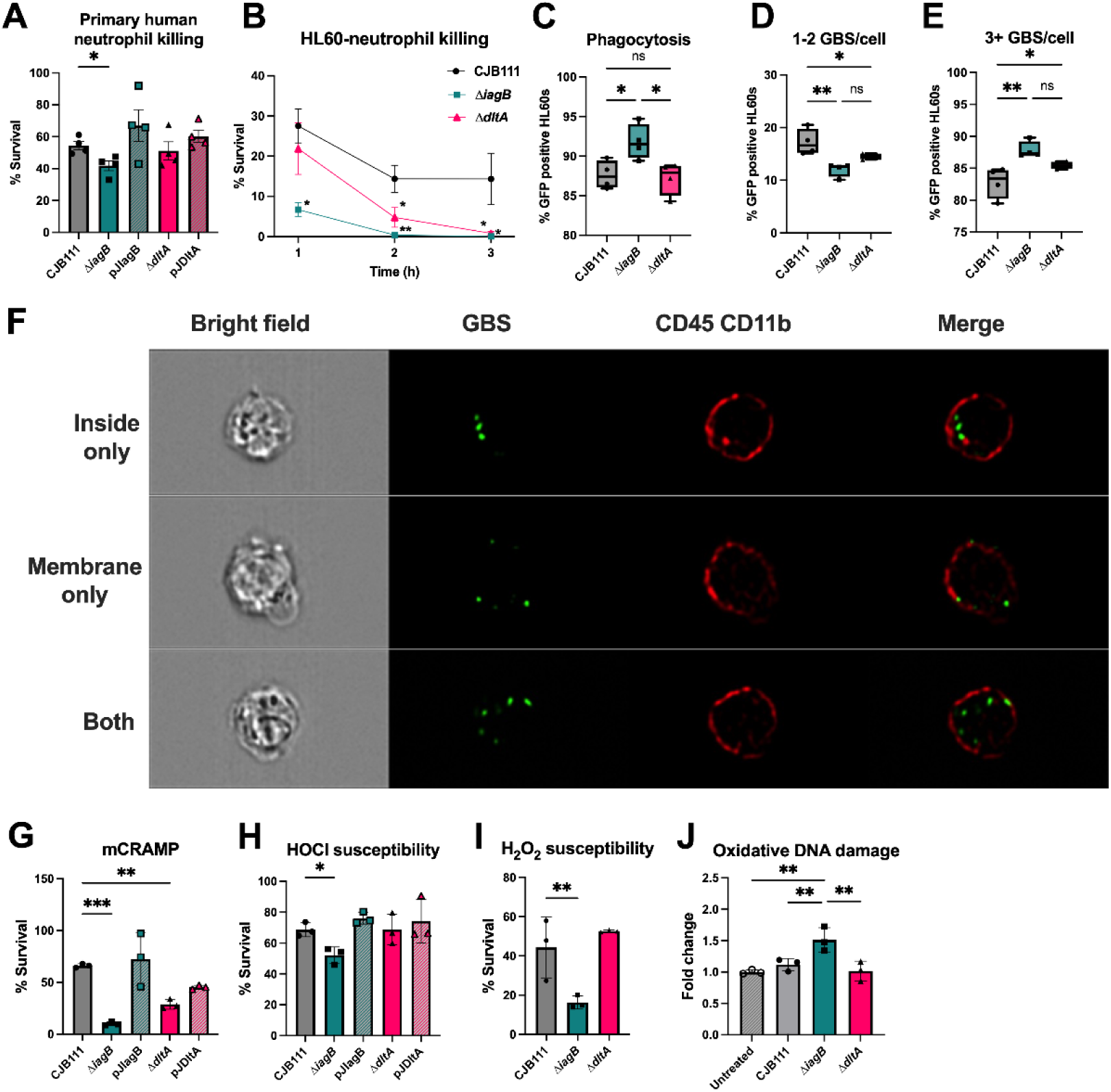
GBS susceptibility to neutrophil killing factors. A) Primary human neutrophil killing of GBS WT, GBSΔ*iagB*, pJIagB, GBSΔ*dltA*, and pJDltA in high glucose infected at MOI 25 for 1 h. % Survival calculated as the ratio of CFU with neutrophils versus CFU without neutrophils. B) HL60-neutrophil like cell killing of GBS WT, Δ*iagB* and Δ*dltA* at MOI 0.001 for 3 h. % Survival calculated as the ratio of CFU in normal serum versus CFU in heat-killed serum in the presence of HL60 cells. C) Phagocytosis frequency of GBS WT, GBSΔ*iagB,* and GBSΔ*dltA* by HL60 cells infected at MOI 8-10 measured by imaging flow cytometry indicated as percent GFP positive HL60 cells, D) 1-2 GBS per cell and E) 3 or more GBS per cell. GBSΔ*iagB* is phagocytized at a higher rate than GBS WT and both GBSΔ*iagB* and GBSΔ*dltA* have higher number of bacteria per neutrophil compared to WT. F) Representative image of GBS WT phagocytosis in HL60 cells. G) Percent survival of GBS WT, GBS*ΔiagB*, pJIagB, GBSΔ*dltA*, and pJDltA to D) mouse cathelicidin mCRAMP after 30 min and H) hypochlorite (HOCl). I) Percent survival of GBS WT, GBS*ΔiagB*, and GBSΔ*dltA*, to 100 µg H_2_O_2_ after 1 h and J) Oxidative DNA damage caused by H_2_O_2_ measured via ELISA. A) Mean and SEM, B,G-J) Mean and SD. C-E) Min to Max. Significance determined by A) Mann-Whitney B-J) One-way ANOVA with Fishers LSD test; *p<.05, **p<.01, ***p<.001. Each assay was performed in at least biological triplicate.

Since GBSΔ*iagB* and GBSΔ*dltA* are more susceptible to neutrophil killing, we next assessed if these mutants exhibited an altered phagocytic uptake compared to WT using imaging flow cytometry. Opsonized GFP-labelled GBS WT, GBSΔ*iagB*, and GBSΔ*dltA* were exposed to HL60 cells for 15 minutes. Interestingly, we observed a significantly higher percent of GFP-positive cells when incubated with GBSΔ*iagB* compared to WT and GBSΔ*dltA* (Figure 4C). We next assessed how many bacteria are present in each cell that was GFP positive (Figure 4D,E) and observed GBS WT had more low numbers (1-2 GBS per cell) compared to the mutant strains, whereas GBSΔ*iagB* and GBSΔ*dltA* had significantly higher numbers (3 or more GBS/cell) compared to WT. Representative images are shown in Figure 4F. Interestingly, over 80% of GFP positive HL60s had GBS both inside and membrane associated compared to the inside or membrane only location, confirming phagocytosis of GBS (Figure S5D-F). These data demonstrate that the loss of GBS glycolipids results in higher frequency in phagocytosis, which along with the loss of LTA alanylation results in increased bacteria per neutrophil.

### Glycolipids contribute to GBS survival against CAMPs and ROS

Neutrophils utilize multiple antimicrobial components to kill bacteria, especially CAMPs such as cathelicidin and ROS such as hydrogen peroxide and hypochlorite ^78–81^. The alanylation of LTA has previously been shown to contribute to GBS evasion of CAMPs by regulating the density of the cell wall ^64^. Similarly, we have recently demonstrated that all three GBS glycolipids aid in the protection against CAMP-mediated killing, likely through a charge repulsion mechanism ^43^. Here, we observed GBS*ΔiagB* is more susceptible to neutrophil killing compared to GBS WT and to GBS*ΔdltA*. To assess if the same is true with CAMP susceptibility, empty vector strains of GBS WT, GBS*ΔiagB*, GBS*ΔdltA*, and complement strains GBS*ΔiagB+*pJIagB and GBS*ΔdltA*+pJDltA were incubated with 20 µM of the mouse cathelicidin, mCRAMP, for 30 min. We observed that the GBS*ΔiagB* mutant exhibited significantly more mCRAMP sensitivity compared to GBS WT and GBS*ΔdltA* strains, and the complemented strains restored survival to WT levels (Figure 4C). Possible explanations for this difference in CAMP susceptibility are alterations in membrane fluidity ^82^. However, no difference in membrane fluidity was observed between all strains as measured by anisotropy, indicating CAMP susceptibility is independent of membrane fluidity in these mutants (Figure S4C).

ROS is another major mechanism of bacterial killing employed by neutrophils and diabetic wounds have increased levels of ROS compared to nondiabetic wounds ^83,84^; thus, we hypothesized that glycolipids and LTA alanylation would contribute to ROS survival. To investigate this, empty vector strains of GBS WT, GBS*ΔiagB*, GBS*ΔdltA*, and complement strains GBS*ΔiagB+*pJIagB and GBS*ΔdltA*+pJDltA were incubated with 0.08% hypochlorite (HOCl) for 1 h resulting in a significant decrease in survival of the GBS*ΔiagB* mutant compared to GBS WT, which was rescued to WT survival levels in the GBS*ΔiagB+*pJIagB complement strain (Figure 4D). No difference in survival was observed between GBS*ΔdltA,* GBS*ΔdltA*+pJDltA, and GBS WT. To further confirm ROS susceptibility and assess oxidative DNA damage; GBS WT, GBSΔ*iagB* and GBSΔ*dltA* were exposed to 100 µg of hydrogen peroxide for 1 h resulting in a significant decrease in survival of the GBS*ΔiagB* mutant compared to GBS WT. No difference between GBS WT and GBSΔ*dltA* was observed (Figure 4E). Similarly, we measured oxidative DNA damage via ELISA and observed a significant increase in fold change of oxidative DNA damage in the GBSΔ*iagB* strain compared to untreated, GBS WT, and GBS*ΔdltA* strains (Figure 4F). No difference was observed between GBSΔ*dltA* and GBS WT or untreated cells. These data suggest that LTA alanylation is important for resisting CAMPs; however, GBS glycolipids promote resistance to both CAMPs and ROS.

### GBS induces primary and secondary degranulation in primary human neutrophils

An important mechanism of neutrophil killing of bacteria is degranulation. Neutrophils produce four types of granules: azurophilic/primary, specific/secondary, gelatinase/tertiary and secretory, with high concentrations of CAMPs, ROS, proteinases and additional antimicrobial factors harbored in the primary and secondary granules ^78–81^. To assess induction of degranulation we exposed primary human neutrophils conditioned in media containing 22 mM glucose to mimic hyperglycemia to opsonized GBS WT, GBSΔ*iagB* and GBSΔ*dltA* for 30 min and assessed degranulation using flow cytometry for established human degranulation markers ^85^ (Figure S3B). We observed a significant increase in azurophilic/primary degranulation, indicated by the increased surface expression of CD63 (Figure 5A) and specific/secondary degranulation, indicated by the increased surface expression of CD66b (Figure 5B) and CD15 (Figure S6B), of neutrophils exposed to GBS regardless of the GBS strain, compared to uninfected control neutrophils. Interestingly, the GBSΔ*iagB* mutant, even though more susceptible to ROS and CAMPs, induces significantly higher levels of azurophilic/primary degranulation compared to GBS WT and GBSΔ*dltA* mutant. While GBS infection resulted in no difference in surface expression of gelatinase/tertiary degranulation marker CD11b between infected and uninfected neutrophils, we did observe an increase in surface CD18 expression on infected neutrophils compared to uninfected neutrophils, suggesting tertiary degranulation is not stimulated or is stimulated to a lesser extent (Figure S6C).

**Figure 5.**
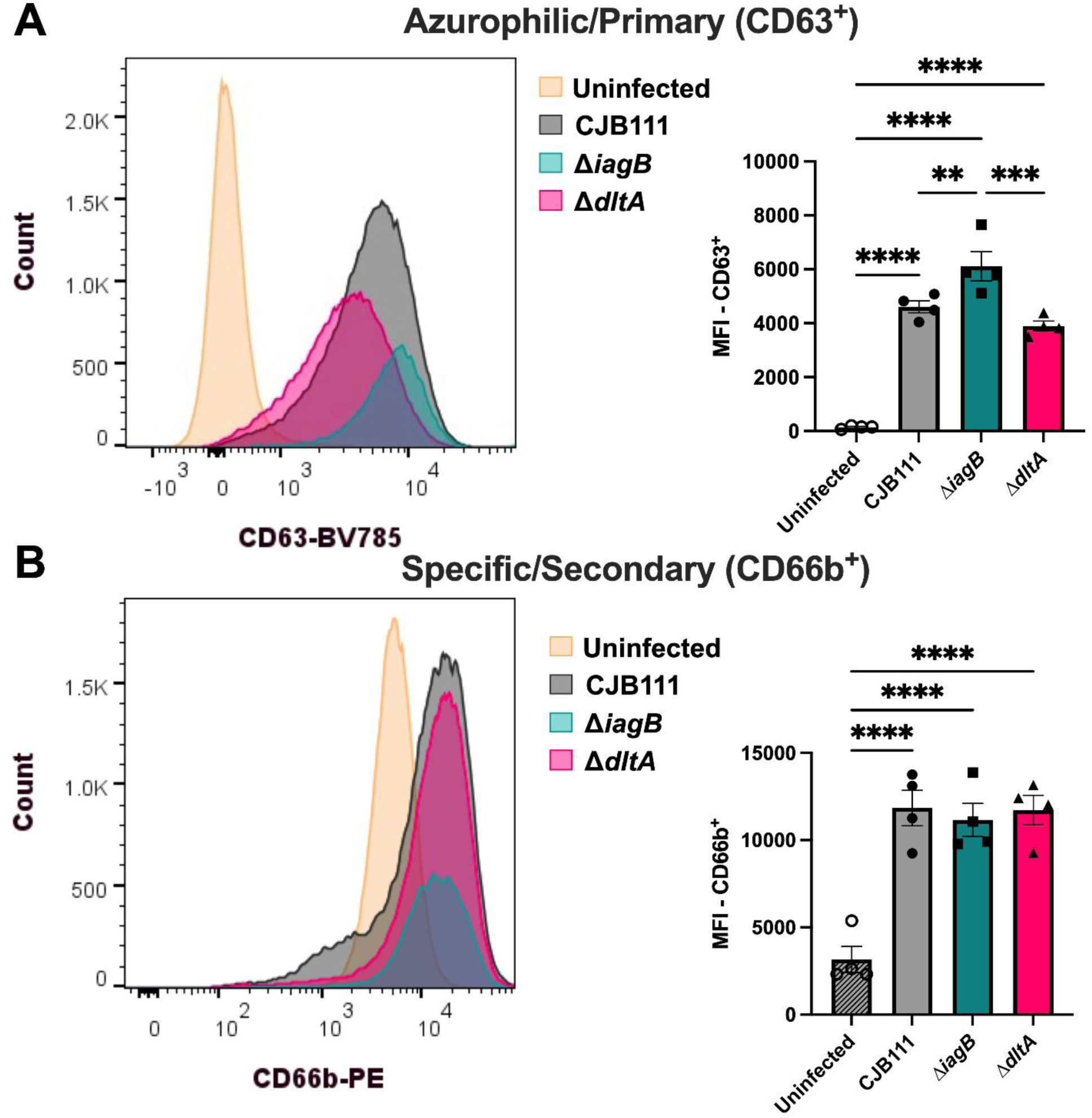
GBS glycolipids and LTA alanylation is not required for degranulation in primary human neutrophils. Neutrophil degranulation measured in primary human neutrophils conditioned in high glucose exposed to opsonized GBS by flow cytometry after 30 min. A) Azurophilic/primary (CD63^+^) granules: left; representative histogram of counts of CD63^+^ neutrophils and right; median fluorescent intensity (MFI) of CD63. Specific/secondary granules B) CD66b^+^. A significant increase in both primary and secondary degranulation compared to uninfected neutrophils is observed. A-C) Mean and SEM. A-C) Significance determined by One-way ANOVA with Fisher’s LSD test; *p<.05, **p<.01, ***p<.001, ****p<.0001, ns; not significant. Assay was performed in at biological quadruplicate.

These data indicate that neutrophils recognize GBS, regardless of the missing glycolipids or LTA alanylations, and trigger primary and secondary degranulation to combat them.

### Neutrophils influence GBS burden in the wound and kidneys of diabetic mice

Neutrophils are highly abundant during diabetic wound infection and promote high inflammation in the microenvironment ^46,86^, as we have also observed during GBS infection (Figure 1). We therefore sought to investigate the contribution of neutrophils to GBS clearance in a diabetic host, and we hypothesized that depletion of neutrophils in the diabetic wound would lead to enhanced recovery of the GBSΔ*iagB* and GBSΔ*dltA* mutants in comparison to non-depleted controls. To investigate this, systemic neutrophil depletion of diabetic mice was performed via intraperitoneal injection of anti-Ly6G antibody, and non-depleted mice were injected with isotype (anti-IgG2a antibody) control one day before infection. Neutrophil depletion in wound tissues was confirmed by flow cytometry (Figure S7A,B). Neutrophil depleted and control diabetic mice were infected with either GBS WT, GBSΔ*iagB* or GBSΔ*dltA* mutants using our streptozotocin induced diabetic wound infection model. Depletion of neutrophils lead to a significant increase bacterial burden of GBS WT and GBSΔ*iagB* as well as an increase in GBSΔ*dltA* in the diabetic wound compared to the non-depleted controls at 48 hpi (Figure 6A). While GBS burdens were consistently higher in depleted compared to non-depleted controls, the GBSΔ*iagB* and GBSΔ*dltA* mutants still exhibited a decreased burden when compared to GBS WT within the neutrophil depleted group (Figure 6A). Similarly, depletion of neutrophils in non-diabetic mice resulted in significantly higher recovered CFU and greater weight loss in the depleted group vs control (Figure S7C,D). Depletion of neutrophils in diabetic mice also led to greater GBS dissemination into the kidneys based on CFU recovered (Figure 6B). Interestingly, significant increases in bacterial burdens in the kidneys of depleted mice infected with GBSΔ*iagB* and GBS*ΔdltA* compared to non-depleted mice was observed and a higher percentage of mice had CFU in the kidneys in the depleted groups vs non-depleted groups (Figure 6C). Taken together these data indicate that neutrophils play an important role in controlling GBS persistence in the wound and preventing systemic spread in the diabetic host. They further suggest that circulating neutrophils can control both the GBSΔ*iagB* and GBS*ΔdltA* during invasive infection *in vivo*.

**Figure 6.**
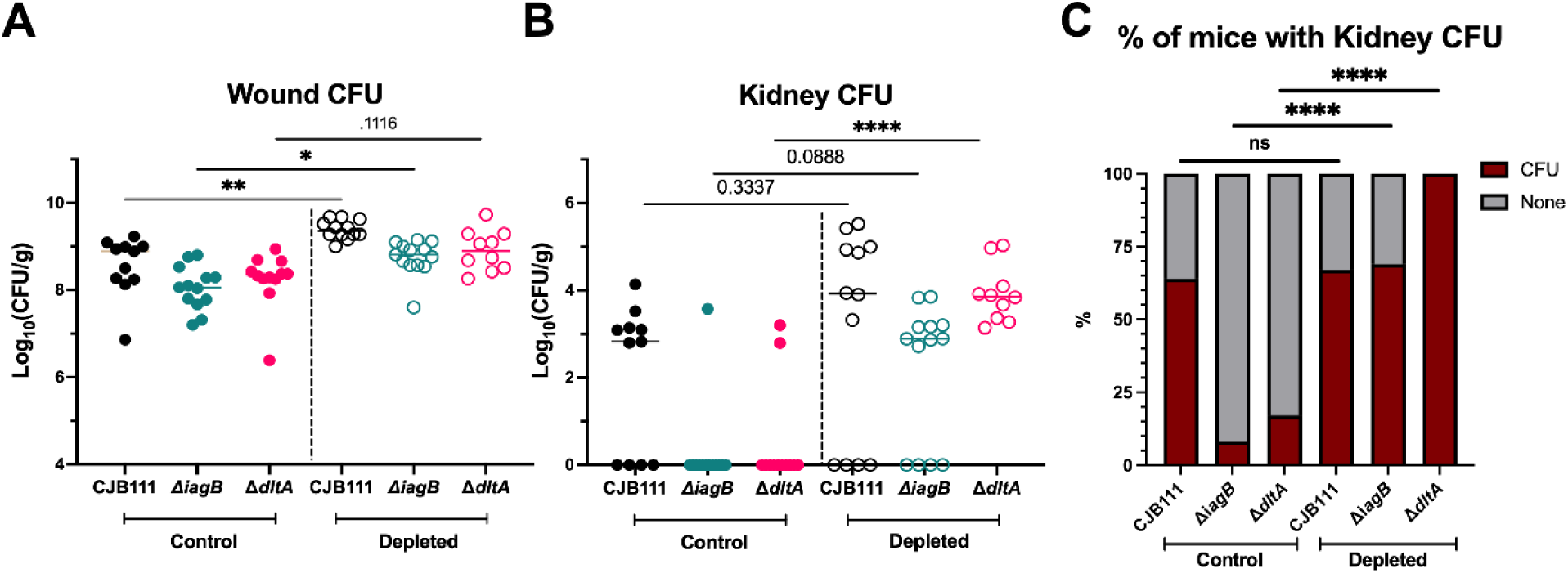
Neutrophils play an important role in combating GBS in the diabetic wound. Control (anti-IgG2a antibody, closed circles) and neutrophil depleted diabetic mice (anti-Ly6G antibody, open circles) were infected with 1 x 10^7^ GBS WT, Δ*iagB*, and Δ*dltA* and bacterial burdens were assessed at 48h in the A) wound and B) kidney. Significantly higher bacterial burdens were recovered in the wound and kidney in depleted mice compared to control mice. C) % of mice in each group with CFU in the kidney. Control: GBS WT; *n =* 11, Δ*iagB; n* = 13, and Δ*dltA;* n *=* 12. Depleted: GBS WT; *n =* 12, Δ*iagB; n* = 13, and Δ*dltA;* n = 10. Significance determined by A,B) Kruskal-Wallis test with Dunn’s multiple comparisons, C) Fishers exact test; *p<.05, **p<.01, ****p<.0001.

## DISCUSSION

The diabetic wound is hyperinflammatory, with high levels of neutrophils and macrophages potentiating inflammation and limiting reformation of the extracellular matrix^87^. However, prior to this work it was unknown how the presence of GBS in diabetic wound infection would affect immune cell populations. Using flow cytometry, we found that neutrophils are the most abundant immune cell GBS encounters during initial diabetic wound infection (48 hpi), making up greater than 65% of all immune cells. We also found a significant increase in the percentage of neutrophils residing in the diabetic wound infected with GBS compared to uninfected controls. We then compared the immune populations in GBS infected wounds in a healthy and a diabetic host. While neutrophils were the most abundant population regardless of host status, we saw a significant decrease in the percentage of live immune cells in diabetic mice, suggesting GBS may be killing immune cells at a greater level in a host with diabetes. In addition to flow cytometry, we performed multispectral immunohistochemistry on GBS and mock infected wounds to visualize the localization of both GBS and immune cells during diabetic wound infection. Wounds infected with GBS led to the formation of a large bolus of bacteria within the wound underneath the area of initial incision. This area contained GBS that seemed to co-localize with neutrophils at the wound site. This was not the case for all immune populations as macrophages were more present in the tissue section from mock infected wounds in comparison to GBS infected. These data are the first to spatially characterize the GBS-host interactions within diabetic wound infection and demonstrate the proximity of neutrophils and the bacterium within the wound. Further, it highlights the need for mechanistic studies that identify GBS factors that contribute to neutrophil survival in this niche.

Despite the growing evidence that GBS is present in diabetic wound infections, little is known about the genetic factors required for fitness and survival in the wound. Here, we used a highly saturated Tn-seq library and identified 291 significantly underrepresented genes in diabetic wound infection, indicating they may be required for fitness. One of the most significantly underrepresented genes was *iagA*, which encodes for an enzyme involved in GBS glycolipid biosynthesis and is responsible for synthesizing the LTA anchor, Glc_2_-DAG, ^43,45^. Additionally, multiple genes encoding enzymes involved in the alanylation of GBS LTA were significantly under-represented ^65^, suggesting that both LTA anchoring and modification are important for GBS to establish itself in this niche. Further investigation of the dataset identified *mprF,* another gene in the GBS glycolipid biosynthesis pathway, and the enzyme responsible for lysinylation of Glc-DAG and phosphatidylglycerol ^44^. Two genes in these biosynthetic pathways (*iagB* and *ltaS*) were not identified in our Tn-seq screen, however, very few transposon insertions in these genes were present in the input library. *LtaS* is known to be essential in other Gram-positive bacteria ^88,89^; however, *iagB* is not essential thus highlighting a limitation of largescale datasets and potential for false negative hits. In this work, we characterize the role of GBS glycolipids and the alanylation of LTA in diabetic wound infection. Using our isogenic glycolipid mutants (Δ*iagB, ΔiagA, ΔmprF*) and our newly constructed LTA alanylation mutant (Δ*dltA*), we confirmed our Tn-seq results observing significant attenuation of these mutants in the diabetic wound compared to GBS WT 48 hpi Throughout this work, we focused on the mechanisms by which all GBS glycolipids and LTA modification contribute to survival in the diabetic wound using the GBSΔ*iagB* (lacking all glycolipids) and GBSΔ*dltA* (unable to add alanine modifications to LTA) mutant strains. Analysis of these mutants at an early time point (5 hpi) revealed that CFU was similar between all strains, demonstrating that these mutants are able to establish infection and are cleared between 5 and 48 h. Further, levels of MPO and CXCL1 were similar in all infected tissues suggesting that immune cells are called to the site of infection at a similar rate. Further we show that complementation of GBS*ΔiagB* in the diabetic wound restores CFU to GBS WT levels (Figure S7E).

Little is known about GBS interaction with keratinocytes, with the only factor known to aid adherence to keratinocytes being the adhesin, PbsP ^41^. Interestingly, we observed the GBS glycolipids and LTA alanylation do not play a role in adherence to human keratinocytes but are important for invasion into the intracellular compartment, as has been observed previously for the BBB endothelial cells ^43–45^. Further, we found that the GBS*ΔiagB* mutant had reduced intracellular survival over two hours of incubation with keratinocytes when compared to WT and other strains. These data suggest that all the glycolipids together contribute to survival once inside keratinocytes. The glycolipid mutant, GBS*ΔmprF,* is more susceptible to low pH compared to GBS WT ^44^ highlighting the glycolipid mutants may be more susceptible to lysosomal killing compared to GBS WT once inside the host cell. It is currently unknown what receptors GBS binds to on keratinocytes, the mechanisms of invasion, and additional GBS factors such as membrane vesicles that may promote invasion into the intracellular compartment of keratinocytes. Thus, further investigation into the mechanisms of interactions of GBS and keratinocytes are warranted, as this may serve as a mechanism to escape from immune cells in the wound and act as potential reservoirs for persistence colonization of the diabetic wound after debridement of the wound.

We have previously found that GBS glycolipids are important for survival against neutrophils, as strains lacking glycolipids have enhanced susceptibility to antimicrobial peptides. However, these studies were performed with neutrophils cultured in normoglycemic levels of glucose. Individuals with diabetes are known to have aberrant neutrophil function, largely driven by hyperglycemia ^90^. We therefore cultured neutrophils in media containing levels of glucose that are physiologically relevant in a diabetic host to assess the role of GBS glycolipids and LTA alanylation to neutrophil survival in high glucose conditions. Interestingly, we observed GBSΔ*iagB* is phagocytized at a higher frequency than GBS WT and GBSΔ*dltA* and both mutants had higher number of GBS per neutrophil compared to GBS WT. These observations demonstrate that GBS glycolipids are important for preventing uptake by neutrophils and once engulfed, important for protecting against killing mechanisms. Here, we also focused on a second mechanism of neutrophil killing, degranulation, using our glycolipid devoid strain, GBS*ΔiagB*, and our LTA alanylation mutant, GBS*ΔdltA*. Using primary human neutrophils exposed to high glucose conditions, we observed all GBS strains, regardless of mutation, triggered both primary and secondary degranulation. This suggests that the mechanism by which neutrophils recognize GBS and elicit degranulation is not dependent on GBS glycolipids or LTA modification. Neutrophil granules, particularly the primary/azurophilic and secondary/specific granules, harbor a variety of antimicrobial properties such as CAMPs and ROS. Due to the hyperglycemia in diabetic individuals, neutrophils are transcriptionally primed to undergo NETosis ^46^. Work by Derré-Bobillot et al. identified a secreted GBS nuclease, NucA involved in the degradation of the DNA matrix, or major backbone of NETs ^46,91^. Further, our group has found that this nuclease is highly upregulated in bacteria recovered from the diabetic wound in comparison to input ^41^. NucA was not identified in the Tn-seq screen, likely because it is a secreted factor that the rest of the population is still producing, preventing it from being underrepresented; however, it may be important in the diabetic wound microenvironment, and its contribution to GBS survival in the diabetic wound is ongoing.

Cationic antimicrobial peptides such as cathelicidin and defensins are small peptides used by the host to target invading pathogens and are an important antimicrobial factor in neutrophil granules. We and others have shown that GBS glycolipids and LTA alanylation are important for resisting CAMP mediated killing although potentially via different mechanisms ^43,64,76^. Saar-Dover *et al* observed that both GBSΔ*dltA* and GBSΔ*mprF* have a more negatively charged cell surface; however, the *mprF* mutant is not susceptible to CAMPs ^76^. Interestingly, they observed the GBSΔ*dltA* has a significantly altered cell wall density resulting in CAMP susceptibility. Based on TEM images here and our previous study, we identified the GBSΔ*iagB* mutant has a significantly more negatively charged cell membrane compared to GBS WT, suggesting the susceptibility to CAMPs is via different mechanism^43^. Interestingly we observed that GBSΔ*iagB* was significantly more attenuated than the GBSΔ*dltA* after 30 min of exposure to mouse cathelicidin, even though the membrane fluidity is the same between these mutants and GBS WT. A possible explanation is that while the alanylation of LTA is required for proper cell wall density to mediate CAMP resistance^76^, the loss of the glycolipids, including the LTA anchor, results in a more severe change in cell wall dynamics possibly making the membrane hyper-attractive to the positively charged CAMPs. Another major antimicrobial property of neutrophil granules is the production of ROS, especially hydrogen peroxide which is used by myeloperoxidase to generate hypochlorite. Not only do neutrophils generate high amounts of ROS, but the diabetic wound contains a large amount of ROS compared to non-diabetic wounds ^83,84^. Like the CAMP phenotype, we observed GBS devoid of glycolipids is more susceptible to both hypochlorite and hydrogen peroxide killing and resulting oxidative DNA damage compared to both WT and Δ*dltA* mutant. This could be due to increased lipid peroxidation of the cellular membrane lacking glycolipids which may generate even more ROS, resulting in increased damage to the cell and death ^92^, but further investigation is needed to identify why glycolipids may protect against ROS attack.

Among the significantly underrepresented genes identified in this Tn-seq are multiple factors required for metabolism such as purine metabolism, glutamine metabolism, and mannose utilization. These genes are important for bacterial pathogenesis ^93^ and were recently identified to be important for *Enterococcus* colonization of non-diabetic wounds ^94^. We also identified multiple GBS virulence factors such as the β-H/C *cyl* operon and the type VII secretion system (T7SS) as under-represented in the diabetic wound. The loss of the GBS β-H/C has been previously shown to impact bloodstream survival and phagocytic defense, and we have also observed its importance to diabetic wound persistence ^41,95^. Similarly, the role of the T7SS has recently been implicated in cytotoxicity to host cells and promotes colonization of the vaginal tract ^59,96^. With the high number of neutrophils and macrophages present in the diabetic wound the T7SS may have important functions for defending against these immune cells and potentially against other bacterial species that colonize this polymicrobial environment. Interestingly, multiple two component systems (TCS) were significantly underrepresented in the Tn-seq. One such TCS, CiaH/R has been implicated in monitoring cell wall/surface stress ^62,97^. GBS TCS 8, which has homology to the VicR/K (WalR/K in some species) system in other streptococci was also significantly underrepresented ^61,98^. Interestingly three currently uncharacterized TCS systems, TCS 7, 12 and 19 ^61^ were also significantly underrepresented. The fact that these TCS systems were significantly underrepresented suggests the strong need for GBS adaptation to the diabetic wound microenvironment. It further suggests these TCS systems may play a pivotal, currently unknown, role in GBS niche adaption, which requires further investigation.

Overall, using Tn-seq, this work has identified multiple GBS factors that are required to colonize the diabetic wound and to evade the large number of immune cells present there (Figure 7). Neutrophils are important first line responder to invading pathogens and the ability for GBS to evade the multiple killing mechanism they possess is vital to establishment in this niche. The characterization of the role that GBS glycolipids and LTA alanylation play during colonization of the wound and evasion of neutrophil killing highlights important cell wall properties that could be targeted by small molecules to aid in the clearance of GBS. Future studies aim to elucidate the roles of other genetic factors identified in the Tn-seq and their contribution to diabetic wound colonization. Ultimately, this work will provide increased knowledge of GBS pathogenesis and virulence potential, aiding in future treatments and may have broad implications to other bacterial species that may colonize the diabetic wound.

**Figure 7.**
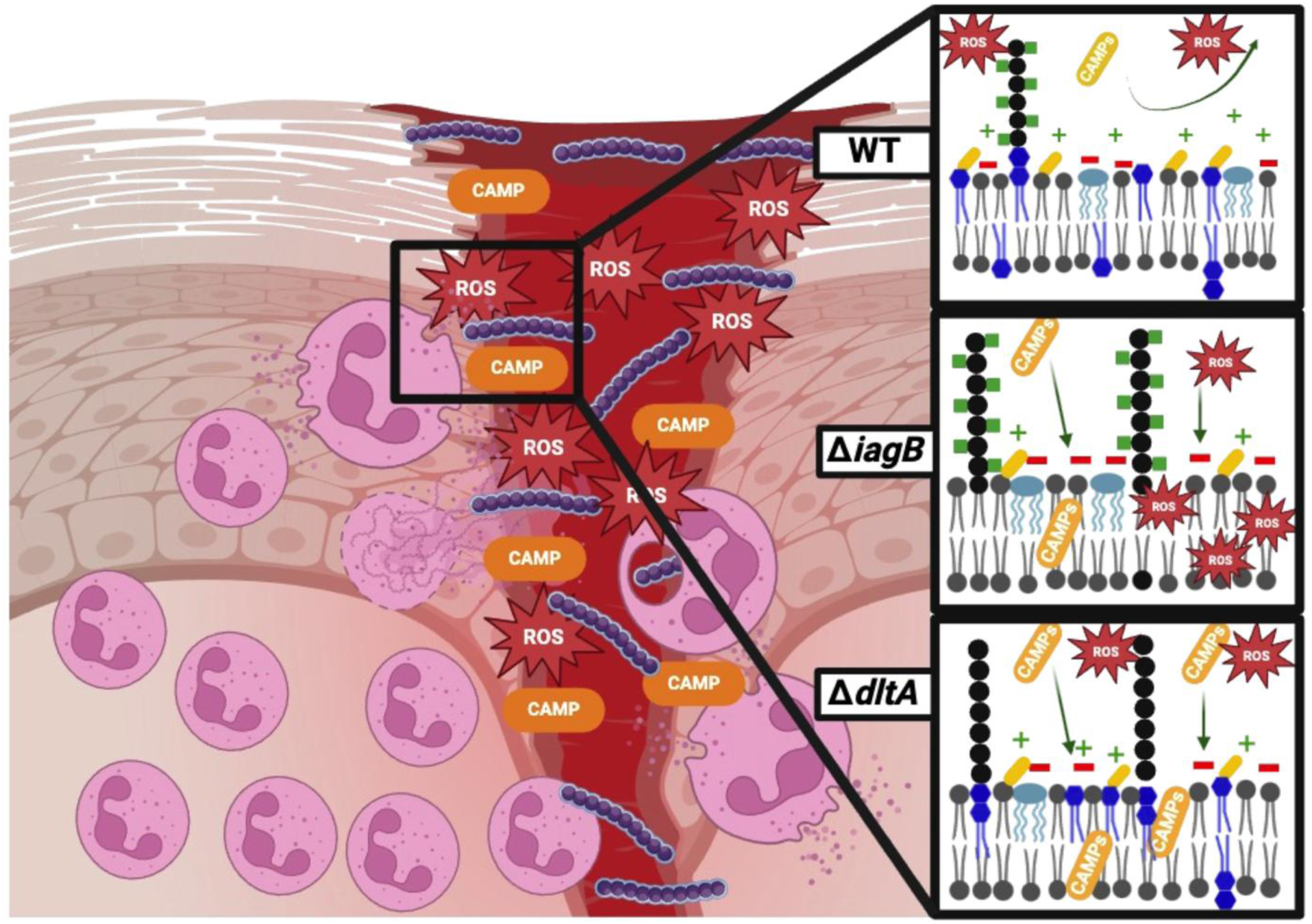
GBS glycolipids and LTA alanylation promote diabetic wound colonization. GBS infects the diabetic wound where it encounters a large population of neutrophils in the wound. Neutrophils are recruited to the wound to control the infection. The diabetic wound is rich in multiple antimicrobial molecules such as cationic antimicrobial peptides (CAMP) and reactive oxygen species (ROS). GBSΔ*dltA* is unable to modify lipoteichoic acid (LTA) and is more susceptible to both neutrophil killing and CAMPs compared to GBS WT, whereas *GBS*Δ*iagB* lacking all glycolipids is hypersusceptible to neutrophil killing, CAMPs and ROS attack.

## ACKNOWLEDGMENTS

The authors would like to thank the members of the Doran and Horswill labs for constructive discussions in the preparation of this manuscript, Dr. Robb Welty at the Biophysics Shared Research Facility at the University of Colorado Anschutz Medical Campus for help with anisotropy experiments, the Electron Microscopy Core Facility at the University of Colorado Anschutz Medical Campus for help with TEM and the Human Immune Monitoring Shared Resource, specifically Troy Schedin and Angela Minic within the University of Colorado Cancer Center (P30CA0434) for their expert assistance in analysis of diabetic wound tissues. This work was funded in part by the American Heart Association grant 23POST1013835 to L.R.J, NIH/NIAID F31AI178881 to M.S.A, F32AI172126 and K99AI180373-02 to R.A.K., R21AI171853 to K.S.D and A.R.H, R01AI178692 to K.S.D.

## AUTHOR CONTRIBUTIONS

Conceptualization: LRJ, KSD, RAK; Methodology: LRJ, MSA, DTN, BLS, RAK; Validation: LRJ, MSA, DTN, BLS, JK, KSD, RAK ; Formal analysis: LRJ, MSA, DTN, JK, RAK; Investigation: LRJ, MSA, DTN, RAK; Resources: JK, KSM, ARH, KSD; Writing original draft: LRJ, RAK; Editing and reviewing: LRJ, MSA, DTN, BLS, JK, KSM, ARH, KSD, RAK; Visualization: LRJ, DTN, RAK; Project administration: LRJ, KSD, RAK; Funding acquisition: LRJ, KSM, ARH, KSD, RAK

## Materials and Methods

### Bacterial strains

All strains used in the study are listed in STAR Methods. GBS strains were grown statically at 37°C in Todd-Hewitt broth (THB). *Escherichia coli* strains were grown in lysogeny broth at 37°C or 30°C with rotation at 225 rpm. Erythromycin and spectinomycin (Sigma-Aldrich) were supplemented to media at 300 and 100 μg/mL for *E. coli* or 5 and 100 μg/mL for GBS strains, respectively.

### Animals

For murine diabetic wound 8 to 12-week-old male mice from the C57Bl/6J or *Lepr^db^* background were used. Animal experiments were approved by the Institutional Animal Care and Use Committee at University of Colorado Anschutz Medical Campus protocol no. 00987 and were performed using accepted veterinary standards. The University of Colorado Anschutz Medical Campus is AAALAC accredited, and its facilities meet and adhere to the standards in the Guide for the Care and Use of Laboratory Animals.

### Routine molecular biology techniques

All PCR reactions utilized Phusion polymerase (New England Biolabs). PCR products and restriction digest products were purified using Qiagen PCR purification kit (Qiagen) per manufacturer protocols. See Table S3 for primers. Plasmids were extracted using Qiagen plasmid mini and maxiprep kits (Qiagen) per manufacturer protocols. Restriction enzyme digests utilized XhoI, PstI, BamHI, and SacII (New England Biolabs) for at least 3 h at 37°C. Ligations utilized T4 DNA ligase (New England Biolabs) at 16°C overnight. All plasmid constructs were sequence confirmed by Nanopore amplicon sequencing (Quintara Biosciences).

### Construction of *dltA* and *cpsD* knockout plasmid

Construction of the *dltA* clean deletion knockout plasmid was performed as previously described ^43,44^. Briefly, approximately 1.5 kb upstream and downstream of the CJB111 *dltA* (ID870_00720), or *cpsD* (ID870_03500) and the spectinomycin cassette from plasmid JC303 ^103^ was amplified using PCR. Plasmid, pMBSacB ^101^, and the PCR products were digested using appropriate restriction enzymes and ligated overnight and transformed into chemically competent *E. coli* MC1061 as described ^43^. Cultures were pelleted using an Eppendorf 5810R centrifuge at 3900 x g for 10 min at room temperature. Plasmid was extracted as described above and sequence verified via Nanopore whole plasmid sequencing (Quintara Biosciences).

### Deletion and complementation of CJB111 *dltA* and *cpsD*

GBS electrocompetent cells were generated as described and double crossover homologous recombination knockout strategy was performed as described previously^43,44^. For *dltA,* after the second cross-over, GBS was incubated in THB + 0.75 M sucrose and 100 µg/mL spectinomycin at 37°C before plating on THB agar supplemented with 100 µg/mL spectinomycin, whereas *cpsD* is a clean deletiona dn no spectinomycin cassette. Colonies were PCR screened for loss of gene using primers outside of the recombination arms and grown in THB at 37°C and stocked. Sequence confirmation of the mutants was done via Nanopore sequencing (Quintara Biosciences). The pJDltA complementation plasmid was constructed as previously described ^43,44^. CJB111 *dltA* gene was amplified, restriction digested and ligated overnight into pDCErm ^104^ plasmid backbone. Ligations were transformed into *E. coli* for maintenance. Complement plasmids sequence confirmed as above and were electroporated into GBS electrocompetent cells prepared as described above.

### Murine model of diabetic wound infection

8 to 12-week-old male mice from the C57Bl/6J or *Lepr^db^* background were used throughout the study. Two weeks prior to infection mice were given low dose injections of Streptozotocin (50mg/kg) (CAS-No: 18883-66-4) dissolved immediately before injection in 100 µL of 50 mM sodium citrate buffer (pH 4.5) for 5 consecutive days, followed by a glucose chaser (250 µL) to prevent hypoglycemia ^77^. Mice then sat for a minimum of one week for blood sugar to normalize before infection. Mice were infected as previously described ^77^. Briefly, the day before infection mice were anesthetized via isoflurane inhalation, shaved and treated with Nair on the back. Tail snips were performed and blood sugar measured via glucometer to ensure all mice had blood glucose levels >200 mg/dl. The following day, mice were weighed and anesthetized via isoflurane inhalation and injected with lidocaine intra-dermally. Mice were wounded with a 6 mm biopsy punch and inoculated with 1 x 10^7^ CFU GBS or a PBS control unless otherwise indicated. Wounds were wrapped with the surgical adhesive Tegaderm for 5 h before adhesive removal and monitoring of animals. After 48 h of infection, mice were sacrificed and wound as well as kidneys were harvested for further analysis. Tissues were placed into 500 µl of sterile PBS in 2.0-ml conical tubes (Thermo Fisher Scientific) along with 1.0-mm-diameter zirconia/silica beads (BioSpec catalog no. 1107911) and homogenized by bead beating two times for 60 s in a BioSpec mini bead beater. Tissue homogenates were plated on GBS CHROMagar [SB282(B)], which allows only for the growth of GBS (in pink) and Enterococcus spp. (in blue). These experiments were approved by the committee on the use and care of animals at the University of Colorado–Anschutz Medical Campus protocol no. 00987. For neutrophil depletion experiments, mice were injected with 200 µg of either anti-Ly6G (BioXCell, clone 18A, catalog BE0075-1) or anti-IgG2a isotype control antibody (BioXCell, clone 2A3, catalog BE0089) diluted in buffered reagent diluent (BioXCell, catalog IP0070) the day before infection.

### Diabetic wound transposon library infection

Triplicate cultures of the pooled CJB111 pKrmit transposon library ^49^ were grown overnight at 37°C in THB with kanamycin at 300 µg/mL and grown to exponential phase without kanamycin before libraries were normalized to ∼1.5 x 10^7^ CFU/10 µL and *Lepr^db^* mice were infected as described above. Input library and wound homogenate was plated in duplicated on CHROMagar Strep B with 300 µg/mL kanamycin and incubated overnight at 37°C to collect recovered transposon mutants. Bacterial growth from spread plates were collected and 4 mice per library pooled together before genomic DNA was extracted using ZymoBiomics DNA miniprep Kit (Zymo Research).

### Transposon library sequencing

Libraries were prepared and sequenced at the University of Minnesota Genomics Center (UMGC) according to https://www.protocols.io/view/transposon-insertion-sequencing-tn-seq-library-pre-rm7vzn6d5vx1/v1. Briefly, genomic DNA was enzymatically fragmented, and adapters added using the NEB Ultra II FS kit (New England Biolabs), and ∼50 ng of fragmented adapted gDNA was used as a template for enrichment by PCR (16 cycles) for the transposon insertions using mariner-specific (TCGTCGGCAGCGTCAGATGTGTATAAGAGACAGCCGGGGACTTATCATCCAACC) and Illumina P7 primers. The enriched PCR products were diluted to 1ng/ul and 10 ul was used as a template for an indexing PCR (9 cycles) using Nextera_R1 (iP5) and Nextera_R2 (iP7) primers. Sequencing was performed using 150- base paired-end format on an Illumina NovaSeq 6000 system to generate ∼40 million reads per library.

### Tn-seq bioinformatics analyses

Bioinformatic analyses were performed as previously described ^50^ Reads were trimmed using Cutadapt (v 4.2) with the following parameters: sequence length with a minimum of 12 bases, removal of flanking “N” bases, reads were trimmed of 3’ “G” bases, and reads were trimmed with the reverse complemented mariner transposon sequence (ACTTATCAGCCAACCTGTTA). TRANSIT (v 3.2.7) ^105^ was used to align trimmed reads to the CJB111 genome (CP063198) for analysis of transposon insertion sites. The Transit PreProcessor processed reads using default parameters with the Sassetti protocol, no primer sequence, and mapped them to the genome using Burrows-Wheeler Alignment (BWA) ^106^. Insertion sites were normalized using the total trimmed reads (TTR) in TRANSIT and analyzed using the site-restricted resampling to compare the insertion counts recovered in wound vs the input library. Resampling was performed using default parameters, with the addition of ignoring TA sites within 5% of the 5’ and 3’ end of the gene. All sequencing reads have been deposited into NCBI SRA under BioProject accession PRJNA1232139.

### N/TERT adherence, invasion and intracellular survival assays

Human immortalized N/TERT-2G keratinocyte cells ^74,107^ were seeded at 1 x 10^5^ cells in keratinocyte serum-free medium (KSFM, Gibco) supplemented with 0.3 mM calcium chloride (PromoCell) in a 24-well tissue culture treated plate and incubated at 5% CO_2_ and 37°C. Assays to determine the total number of bacteria adhered to host cells or intracellular bacteria were performed using GBS empty vector and complement strain as described previously ^43,44^. Briefly, exponential phase bacteria were normalized to 1 x 10^8^ CFU/mL in PBS and 1 x 10^5^ CFU per well at multiplicity of infection (MOI) of 1. The total cell associated GBS were recovered after 30 min incubation. Cells were washed four times with 1X PBS (Sigma) and detached by addition of 100 μL of 0.25% trypsin-EDTA solution (Gibco) and incubation for 5 min at 37°C before lysing the eukaryotic cells with the addition of 400 μL of 0.025% Triton X-100 (Sigma) and vigorous pipetting. The lysates were serially diluted and plated on THB agar and incubated overnight to determine CFU. Bacterial invasion assays were performed as described above except infection plates were incubated for 2 h before incubation with 100 μg gentamicin (Sigma) and 5 μg penicillin (Sigma) supplemented media for an additional 2 h to kill all extracellular bacteria, prior to being trypsinized, lysed, and plated as described. Experiments were performed in biological triplicate with three technical replicates per experiment.

### Dissociation of wound tissue and blood for flow cytometry

Single cell suspension preparation was performed using methods previously described by our group ^47^. In brief, following euthanasia, mice were perfused with PBS+heparin and wounds were excised, minced with a razor blade, and suspended in 2.4 mL RPMI 1640 medium (Corning). Digestion enzymes were obtained from Miltenyi Biotec Multi Tissue Dissociation Kit 1 and resuspended according to manufacturer instructions. 100 μL Enzyme D, 50 μL Enzyme R, 12.5 μL Enzyme A were added to each tissue sample in RPMI. Tissues were mechanically dissociated using gentleMACS™ C Tubes (Miltenyi Biotec; protocol Multi H) for 30 sec, incubated on a rocking shaker at 37ᵒC for 30 min, followed by an additional 30 sec of mechanical dissociation and 30 min incubation at 37°C while rocking. The digestion mixture was strained through a 70 μm filter, and the single cells were washed with RPMI and spun down at 300 x g for 5 min. The cell pellet was resuspended in Ammonium-Chloride-Potassium red blood cell lysis buffer (150 mM NH4Cl, 10 mM KHCO3, 0.1 mM Na2EDTA) for 1 min at room temperature and diluted in RPMI. Following centrifugation, the cell pellet was resuspended in MACS staining buffer (1X PBS, 2 mM EDTA, 0.5% BSA) to generate single cell suspensions. Blood was obtained from right ventricle cardiac puncture. Approximately 200 μL were taken from each animal and mixed a small amount of heparin suspended in PBS. Ammonium-Chloride-Potassium red blood cell lysis buffer was added to blood samples to a final volume of 1 mL, and incubated for 3 min. Samples were then spun down at 300 x g for 5 min and the resulting pellet was washed again in red blood cell lysis buffer with a repeat spin. The cell pellet was resuspended in MACS staining buffer to generate single cell suspensions as noted above.

### Antibody staining of single cell suspensions for flow cytometry analysis

Antibody staining methods and panels were performed as previously described ^47^. Fixable Viability Dye eFluor™ 506 (ThermoFisher; 1:800 dilution) in PBS was added to single cell suspensions and allowed to incubate for 15 min at room temperature. Conjugated antibodies listed in Table S4 were then added to wound and blood single cell suspensions along with anti-mouse CD16/32 (ThermoFisher, Fc block, clone 93) for 30 min at room temperature. To analyze human neutrophils, antibodies against surface markers listed in Table S4 were added with human FcR Blocking Reagent (Miltenyi Biotec). Stained neutrophils were then washed and fixed using Foxp3/Transcription Factor Staining Buffer Set (eBioscience) according to the manufacturer’s protocol. Cells were analyzed on the LSRFortessa (BD Biosciences) flow cytometer. Data were analyzed with BD FlowJo software version 10.10.0. Single-stained beads (Versacomp Antibody Capture Bead Kit; Beckman Coulter) were used as compensation controls. A detailed gating strategy is included in Figure S1 for wound and blood and Figure S6 for neutrophil degranulation.

### Multispectral Immunohistochemistry and Image Analysis

Diabetic mice were wounded and infected with either WT CJB111 expressing GFP or mock infected with PBS as described above. After 48h of infection, animals were sacrificed and wounds excised and sent to the Human Immune Monitoring Shared Resource (HIMSR) at the University of Colorado Anschutz Medical Center. Multispectral immunohistochemistry (mIHC) analyses were performed using Vectra Automated Quantitative Pathology Systems (Akoya Biosciences) as described previously ^108^. Tissues were formalin-fixed, paraffin embedded and sectioned onto slides. Whole tissues were analyzed with the following antibodies; MsCD11b (Novus, NB110-89474), Msly6G (CST, 87048S), HuCitH3 (abcam, ab5103), MsGFP/YFP (CST, 2956S), msCD11c (CST, 97585S), MsCD45 (CST, 70257S), MsEpCAM (CST, 93790S), MsF480 (CST, 30325S), and DAPI. All slides were deidentified and imaged by the Anschutz Medical Campus Human Immune Monitoring Shared Resource (HIMSR) core (RRID:SCR_021985) on Akoya Biosciences PhenoImager HT scanner. Regions of interest (ROI) were selected, and multispectral images were collected with the 20× objective. A training set of nine representative images was used to train the analyses algorithms for tissue and cell segmentation and phenotyping using in Form software (Akoya Biosciences). Representative autofluorescence was measured on an unstained control slide and subtracted from study slides.

### Primary human neutrophil killing

Primary human neutrophils were isolated from healthy human donors (female *n =* 2, male *n* = 2) per the COMIRB protocol number 17-1926 as previously described ^109–111^. Neutrophils were resuspended in RPMI (Gibco, no phenol red) with 22 mM glucose to mimic diabetes ^46^. Mid-exponential phase GBS were normalized to 1 x 10^8^ CFU/mL in Hanks’ Balanced Salt Solution (HBSS, Gibco), and 1mL GBS was opsonized in 10% pooled human serum (Complement Tech) at 37°C with rotation for 30 min. Opsonized GBS were washed once with HBSS before being resuspended in 1 mL of 4 x 10^6^ neutrophils (MOI of 25) and split into two Eppendorf tubes containing 500 µL each. One tube was incubated at 37°C, with rotation, for 1 h and the second tube used for time 0 h counts. At each time point, 500 µL of 0.05% Saponin (Sigma) was added to the tube and vortex vigorously to lyse neutrophils. 20 µL was removed in duplicate and serial diluted and 50 µL was spread plate on THA containing plates and incubate at 37°C overnight for CFU enumeration. Percent survival was calculated as the ratio of CFU at 1 h verses 0 h and normalized to the no neutrophil control (RPMI only tubes) set to 100%.

### HL60 opsonophagocytic killing

HL60 opsonophagocytic killing assays were performed as previously described ^43^. Briefly, HL60 cells purchased from ATCC (ATCC CCL-240) were differentiated with 1.25% DMSO (Sigma) in RPMI for 4 days. Differentiated HL60 cells were harvested and checked for > 65% viability before starting the assay in accordance with a well-characterized protocol. In a 96 well plate (Corning), 50 µL containing ∼1 x10^3^ CFU of mid-exponential phase GBS, normalized in HBC buffer (1X HBSS, 0.5% BSA, and 2.2 mM CaCl_2_, pH 7.2) was added. 50 µL of either 20% normal human serum (Complement Tech) or 20% heat-killed human serum (60°C for >30 min) in HBC buffer were added to each well and incubated at 37°C, with shaking, for 15 min. 100 µL of HL60 cells (∼5 x10^5^) were added to each well and incubated at 37°C with shaking, MOI ∼0.001, as previously described. Over a 3h time course, 10 µL’s were removed and indicated times, serial diluted and plated on THB agar for enumeration. Assays were performed in biological triplicate with technical duplicates per replicate. Percent survival was calculated as the ratio of CFU in normal serum versus CFU in heat-killed serum, and no HL60 control wells were set to 100%.

### HL60 opsonophagocytic uptake measured by imagestream flow cytometry

GBS were opsonized as described above and HL60-neutrophil like cells were infected at an MOI of 8 - 10 for 15 minutes at 37°C with rotation. Following incubation tubes were transferred to ice and washed in MACS staining buffer described above. To stain the host cell surface, HL-60s were incubated with anti-human CD11b (cat no. 301309, clone [ICRF44]) and anti-human CD45 (cat no. 304011, clone [HI30]) both conjugated to APC along with human FcR Blocking Reagent (Miltenyi Biotec). Cells were fixed using Foxp3/Transcription Factor Staining Buffer Set (eBioscience) according to the manufacturer’s protocol. Samples were run on the Imagestream Mark II cytometer (Cytek Bioscieces) using the 60x objective. Samples were then analyzed with the IDEAS v6.4 software (Cytek Biosciences). For analysis masks were made using the object and adaptive erode functions for generating cell, inside the cell and the membrane masks. The samples were first gated on area vs aspect ratio for single cells, and only in focused cells were included. Focused cells were then gated on CD45^+^CD11b^+^ expression using the membrane mask and intensity per area feature. GBS bacteria on the membrane, inside the cell or both were numerate with the threshold function mask and the spot count feature. The assay was performed in biological quadruplicate.

### Primary human neutrophil degranulation

To measure degranulation in primary human neutrophils, neutrophils were isolated as described above and infected with opsonized GBS at an MOI of 25 in 22 mM glucose as described above for 30 min at 37°C in a 96 well plate. Antibody staining and flow cytometry analysis is described above using known markers of degranulation ^85,112^.

### Cationic antimicrobial peptide killing

mCRAMP killing assays were performed with concentrations as previously described ^43^. A total of 2-4 x 10^5^ CFU of mid-exponential phase cultures were exposed to 20 µM mCRAMP (AnaSpec) in THB for 30 min at 37°C. Dilutions were plated for CFU enumeration.

### Reactive oxygen species susceptibility and 8-OHdG ELISA

Overnight cultures were normalized to 1 x 10^8^ CFU/mL and 2 mL of bacteria were exposed to 0.08% hypochlorite (Clorox) or 100 µg of hydrogen peroxide (Millipore, 386790) for 1 h at 37°C before 10 µL was removed for serial dilution and plating for CFU on THA plates. The rest of the bacteria exposed to hydrogen peroxide were pelleted and stored at -20 °C for 8-OHdG Quantification via OxiSelect Oxidative DNA Damage ELISA Kit (Cell Biolabs, STA-320) per manufacturer protocol. Biological triplicate was performed.

### Transmission electron microscopy

TEM was performed at the electron microscopy core facility at the University of Colorado Anschutz Medical Campus. Approximately 8 mL of mid-exponential (OD_600nm_ 0.5 – 0.6) growing bacteria were pelleted and cells were fixed with 2% paraformaldehyde and 2% glutaraldehyde in 0.1M sodium cacodylate buffer and embedded in 3% agarose. Samples were trimmed into small bocks and rinsed three rinses in 0.1 M sodium cacodylate buffer, they were post-fixed in 2% osmium tetroxide and 0.8 % K_3_[Fe(CN_6_)] in 0.1 M sodium cacodylate buffer for 1 h at room temperature. Cells were rinsed with water and en bloc stained with 2% aqueous uranyl acetate for 1 h. They were dehydrated with increasing concentration of ethanol, infiltrated with SPURR resin and polymerized in a 60^°^C oven overnight. Blocks were sectioned with a diamond knife (Diatome) on a UC7 ultramicrotome (Leica) and collected onto copper grids, post stained with 2% aqueous uranyl acetate and lead citrate. Images were acquired on a Tecnai T12 transmission electron microscope (ThermoFisher) equipped with a LaB_6_ source at 80 kV using an NS15 (15 Mpix) camera (AMT).

### GBS capsule flow cytometry

Flow cytometry for capsule expression was performed using CJB111 WT, CJB111Δ*iagB*, CJB111Δ*iagA*, CJB111Δ*mprF,* CJB111Δ*dltA,* and CJB111Δ*cpsD* empty vector strains, the respective complement strains, pJIagB, pJIagA, pJMprF, and pJDltA, and GBS A909 (Ia Capsule). Overnight cultures were washed in HBC buffer (Hanks’ balanced salt solution [HBSS]-bovine serum albumin-calcium buffer; 1× HBSS without magnesium or calcium, 0.5% bovine serum albumin, 2.2 mM CaCl_2_) and incubated with adsorbed anti-Group B serotype V antisera (SSI Diagnostica Group, cat. no 22461) at a 1:5000 dilution. Bacteria were washed via centrifugation and labeled with a goat anti-rabbit Ig conjugated to AF647 (SouthernBiotech, cat. no 4010-31) at a 1:5000 dilution. Bacteria were fixed in 5% formalin and analyzed on a BD LSRFortessa (BD Biosciences) using the BD FacsDiva software (v9). Data were anlyzed with BD FlowJo software v 10.10.

### Anisotropy

The small molecule probe 1,6-diphenyl-1,3,5-hexatriene (Sigma; DPH) was used to evaluate membrane fluidity as previously described ^113,114^. Briefly, pellets from overnight cultures were washed with phosphate buffered saline (PBS) twice and resuspended in fresh PBS to ∼ 1 x 10^8^ CFU/mL in 3 mL of PBS and incubated with 5 μM of DPH for 15 min at 37°C and 200 rpm and samples were kept at 37°C using a water bath until analyzed. Anisotropy measurements were taken on a Jobin Yvon FluoroMax-3 spectrofluorometer with polarizers in the L geometry at the Biophysics Shared Research Facility at the University of Colorado Anschutz Medical Campus. DPH was excited at 360 nm and the fluorescence intensity was monitored at 430 nm. Samples were maintained at 37°C using the temperature-controlled sample compartment of the spectrofluorometer. Measurements were performed on biological quadruplicate. The anisotropy of DPH was calculated from the differences in the fluorescence intensity detected with the polarizers in the vertical and horizontal orientations and data is reported in arbitrary units (AU). Biological quadruplicate was performed.

## QUANTIFICATION AND STATISTICAL ANALYSIS

Statistical details of each experiment including statistical tests, number of samples, and significance threshold can be found in the figure legends. All statistical analysis was performed using GraphPad Prism software version 10 (San Diego, California) unless otherwise indicated, and statistical significance was accepted at p-values of < 0.05.

